# A global viromic survey reveals unprecedented viral diversity within ammonium-oxidizing archaea and related lineages

**DOI:** 10.1101/2025.08.23.671958

**Authors:** Kaiyang Zheng, Yantao Liang, Fabai Wu, Hao Yu, David Paez-Espino, Mario López-Pérez, Sungeun Lee, Graeme W. Nicol, Christina Hazard, Andrew McMinn, Min Wang

## Abstract

*Nitrososphaeria*, an archaeal class containing all ammonium-oxidizing archaea (AOA) and their closely related clades (e.g. *Ca. Caldarchaeales*), is a widely distributed archaeal lineage across diverse ecosystems. Due to the difficulty of isolation and cultivation, the viral diversity and evolution associated with non-marine AOA lineages and *Ca. Caldarchaeales* remain largely uncharacterized. Here, based on globally sampled genomes and metagenomes, we assembled a genomic catalogue associated with AOA, including 283 species-level viruses and 415 species-level mobile genetic elements (MGEs). Notably, 259 previously unrecognized species-level viruses or MGEs associated with *Ca. Caldarchaeales* were identified. Across the two major *Nitrososphaeria* lineages, we uncovered an expansion of archaeal viral families: 11 families associated with AOA, including seven new head-tailed lineages, one of which comprises a mega archaeal virus with a genome size exceeding 220 kb; ten families associated with *Ca. Caldarchaeales*, including four novel head-tailed groups and two previously unknown rod-shaped family (*Demeterviridae* and *Gaiaviridae*). Phylogenetic inference showed that rod-shaped viruses associated with *Ca. Caldarchaeales* and methanogens are phylogenetically intertwined and likely share a common ancestor with *Hoswirudivirus*, a *Sulfolobales*-associated rod-shaped virus characterized by two copies of the coat gene. Through structural modelling, a rare fusion of the coat protein in *Demeterviridae* was observed, in which a single open reading frame within a gene island adjacent to a glycosyl transferase appears capable of forming a complete capsomer. This provides new insights into the origin and evolution of *Adnaviria* viral realm. Functional genomics analyses further revealed that these novel rod-shaped viruses encode virus-specific rare quinone biosynthesis genes and arsenic response transcriptional regulator genes, providing potential clues to their mechanisms of adaptation to hyperthermal environments. Collectively, our work uncovers the hidden genomic, evolution and functional diversity of *Nitrososphaeria* viromes for the first time and closes a long-standing gap in archaeal virome research.

**Highlights:**

1. First genomic catalogue of viruses and MGEs linked to *Nitrososphaeria*.
2. Discovery of diverse novel archaeal viral families.
3. New insights into rod-shaped virus evolution and archaeal virus–host adaptation.

## Introduction

The *Nitrososphaeria* is an archaeal class within *Thermoprotei*, encompassing the previously defined *Thaumarchaeota* and *Augarchaeota*, both of which are important members of the *Thermoproteati* (previously known as TACK superphylum)^1,2^. In many aquatic ecosystems, *Nitrososphaeria* constitute up to 40% of microorganisms, contributing significantly to ammonia oxidation, thus heavily influencing nitrogen cycling ^3–5^. *Nitrososphaeria* includes all ammonia-oxidizing archaea (AOA) ^6^ and utilizes ammonia oxidation pathways as their primary energy source ^7,8^. Compared to ammonia-oxidizing bacteria, AOA thrive in low-ammonia conditions due to higher ammonium affinity, dominating nitrogen-limited environments as efficient “nitrogen thieves” ^9–14^. The ancestors of AOA are considered to have originated in terrestrial environments and to have adapted aerobically after the Great Oxygenation Event ^13,15,16^. Subsequently, these archaea became key microbial players across diverse ecosystems and make a substantial contribution to carbon fixation and ammonia oxidation ^8,15,17–21^. As the sister clade of AOA, *Ca. Caldarchaeales* shares substantial genetic content involved in sulphur oxidation and ammonia oxidation, suggesting an ancestral ecological transition of AOA from hyperthermal to moderate environments ^22^. Primarily inhabiting hydrothermal vents and hot springs, *Ca. Caldarchaeales* influences carbon and sulphur cycling, nutrient transformation and microbial community structures, playing critical ecological roles in extreme environments ^23,24^.

Viruses and mobile genetic elements (MGEs) exhibit complex morphological and genomic diversity, significantly shaping archaeal genomes and their ecological footprint ^25,26^. Due to cultivation challenges, environmental metagenomics has become the primary method for exploring archaeal viral diversity. Recent studies using environmental metagenomes have greatly expanded viral diversity associated with the Asgard super-phylum, methanogens and DPANN super-phylum, addressing gaps in viral evolution and host-virus interactions ^27–31^. Over 100 AOA-associated viral or MGE sequences from marine or soil samples have so far been identified from metagenomic survey, revealing diverse viral elements that influence ammonia oxidation and host genomic organization ^32–40^. However, these case studies were all based on samples from single geographic regions rather than systematic global surveys, which prevents a comprehensive profiling of the diversity, evolution, and functions of these archaeal viruses. The absence of viruses associated with extremophilic AOA lineages (e.g., *Nitrosocaldales* and *Nitrosotalea*) poses a challenge for resolving phylogenetic placements of AOA viruses within the virosphere. In particular, as a phylogenetic sister clade closely related to AOA and one of the main lineages constituting the TACK super-phylum, *Caldarchaeales*-associated virome remain underexplored and represent a major enigma within virology.

To address this knowledge gap, we systematically explored *Nitrososphaeria*-associated viruses or MGEs diversity based on extensive global genomic and metagenomic datasets. Unlike previous studies that focused on AOA-associated viruses from only single geographic regions^32–36,39,40^, our comprehensive analysis of the *Nitrososphaeria*-associated virome fills a critical gap in the understanding of its genomics, phylogenetics, and functional repertoire. Through the establishment of a genomic catalogue comprising hundreds of viral and MGE sequences, we identified nine previously unrecognized archaeal viral families, including two group of rod-shaped viruses associated with *Ca. Caldarchaeales* that exhibit unique capsomer features and metabolic potential. These culture-independent virus–archaea associations remain to be empirically verified as infections, yet they offer a testable data foundation for subsequent establishment of *Nitrososphaeria*–virus culture systems, thereby providing new insights into the evolution of archaeal virosphere.

## Results

### Overview of the genomic catalogue of *Nitrososphaeria*-associated viruses or MGEs

*Nitrososphaeria* genome database containing 2,137 genomes and metagenome-assembled genomes (MAGs) was constructed from NCBI GenBank and IMG/M databases (Supplementary Fig. 1 and Supplementary Note 1)^41,42^. The virus or MGE database comprised over 15 million contigs from IMG/VR v.4, together with 170 metagenome-assembled contigs originating from hyperthermal, abyssal/hadal ocean, hypersaline, frozen, and acidic environments, including 17 from our published and unpublished studies (Supplementary Dataset 1) ^41,43–46^. The integration of viral and MGE sequences from our pipeline and previous studies enabled the establishment of a genomic catalogue comprising 820 *Nitrososphaeria*-associated viruses or MGEs (Supplementary Fig. 1 and Dataset 2). These were grouped into 698 species-level operative taxonomic units (vOTUs), comprising 286 vOTUs with and 407 MGE OTUs without known viral capsid proteins. In addition, 13 vOTU and 27 MGE OTUs associated with AOA and 103 vOTUs and 151 MGE OTUs associated with *Ca. Caldarchaeales* were newly identified, respectively.

*Nitrososphaeria*-associated viruses or MGEs are widely distributed from tropical to polar regions and across terrestrial to marine environments (Fig. 1a). Most of these regions have not been covered by previous studies focusing on AOA-associated viruses^32–36,39,40^. These viruses or MGEs predominately originated from geothermal, marine, soil and hydrothermal environments, aligning with habitats of AOA, deeply-branched AOA-related lineage and *Ca. Caldarchaeales* (Fig. 1b) ^7,20,22,23,47–50^. This genome catalogue revealed novel viruses associated with nine previously unrecognized archaeal lineages within *Nitrososphaeria* (Fig. 1c), including *Ca. Caldarchaeales*, *Nitrosocaldales*, *Nitrosotalea*, and other deeply branched AOA-related lineages (also known as non-AO *Thaumarchaeota*), thereby expanding the number of lineages known to host these viruses from two to eleven ^32–38,51^.

**Figure 1.**
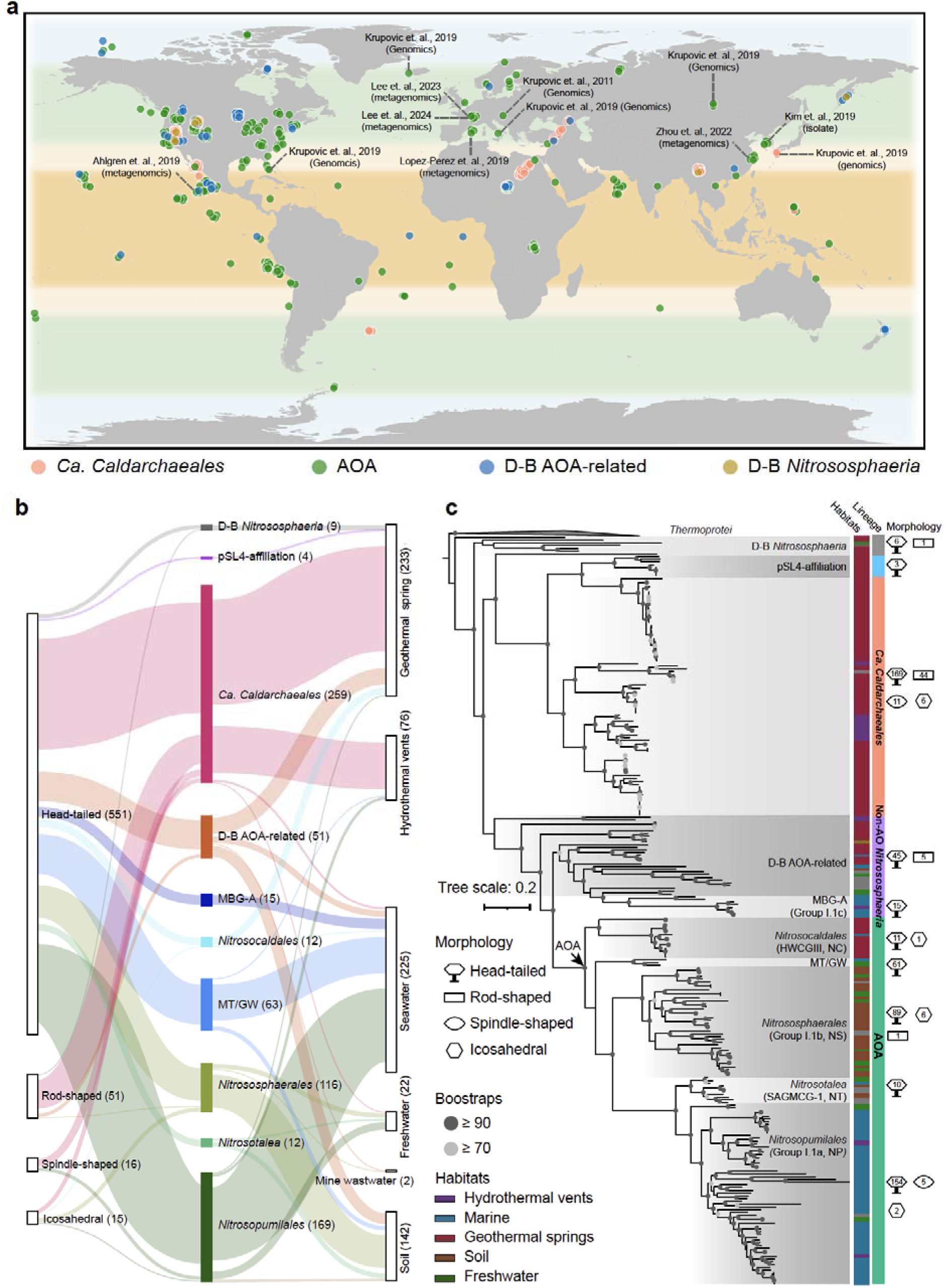
The expanding diversity of *Nitrososphaeria*-associated viruses or mobile genetic elements (MGEs). a, Biogeographic distribution of *Nitrososphaeria*-associated viral or MGE genomes. In the map, nodes of different colors indicate their putative archaeal hosts: *Ca. Caldarchaeales*, ammonia-oxidizing archaea (AOA), deeply-branched AOA-related archaea (DB-AO-related), and deeply-branched *Nitrososphaeria* archaea (D-B *Nitrososphaeria*). Sequences generated or reported in previous studies are indicated alongside the corresponding nodes. b, The number of vOTUs or MGE OTUs with distinct putative morphologies were originated from different habitats and were associated with diverse *Nitrososphaeria* lineages. Bar colours represent different archaeal lineages. c, Phylogenetic tree of *Nitrososphaeria* genomes. This tree was constructed based on concatenated alignments of 46 ribosomal proteins (totalling 5,963 amino acid positions) using the maximum likelihood method with the LG+C60+F model as recommended by Spang et al. and Alves et al. ^2,112^. Reference genomes from *Thermoprotei* were used as the outgroup to root the tree. Tree annotation includes the habitats and lineages of these archaeal genomes. Virion icons to the right of the tree indicate the numbers of viruses or MGEs with different possible morphologies (at the OTU level) associated with each archaeal lineage.

### The expansion of *Nitrososphaeria* virome diversity revealed numerous previously unrecognized archaeal viral families

*Nitrososphaeria*-associated viruses or MGEs exhibit distinct differences in genome content compared with other viruses. The genome similarity network (sharing at least 20% of protein clusters) yielded 67 *Nitrososphaeria*-associated viral communities (super viral clusters, sVCs), with 90% of these communities containing only *Nitrososphaeria*-associated viruses or MGEs (Fig. 2). AOA-associated sVCs and *Caldarchaeales*-associated sVCs show no overlap, indicating clear differences between the viromes of these two main *Nitrososphaeria* lineages. Notably, both exhibit overlaps with the sVCs associated with non-AO *Nitrososphaeria*, which represents a transitional group (n = 6), suggesting the possible presence of transitional lineages within the *Nitrososphaeria*-associated viromes. In addition, some *Nitrososphaeria*-associated viruses exhibited close genome-content relationships with other archaeal or bacterial viruses (e.g. ANME-1 and *Bacteroidota*), suggesting that these *Nitrososphaeria*-associated viruses or MGEs may have undergone host switching or engaged in gene exchange with viruses of other prokaryotes. (Figs. 2 and Supplementary Dataset 3) ^29,30^.

**Figure 2.**
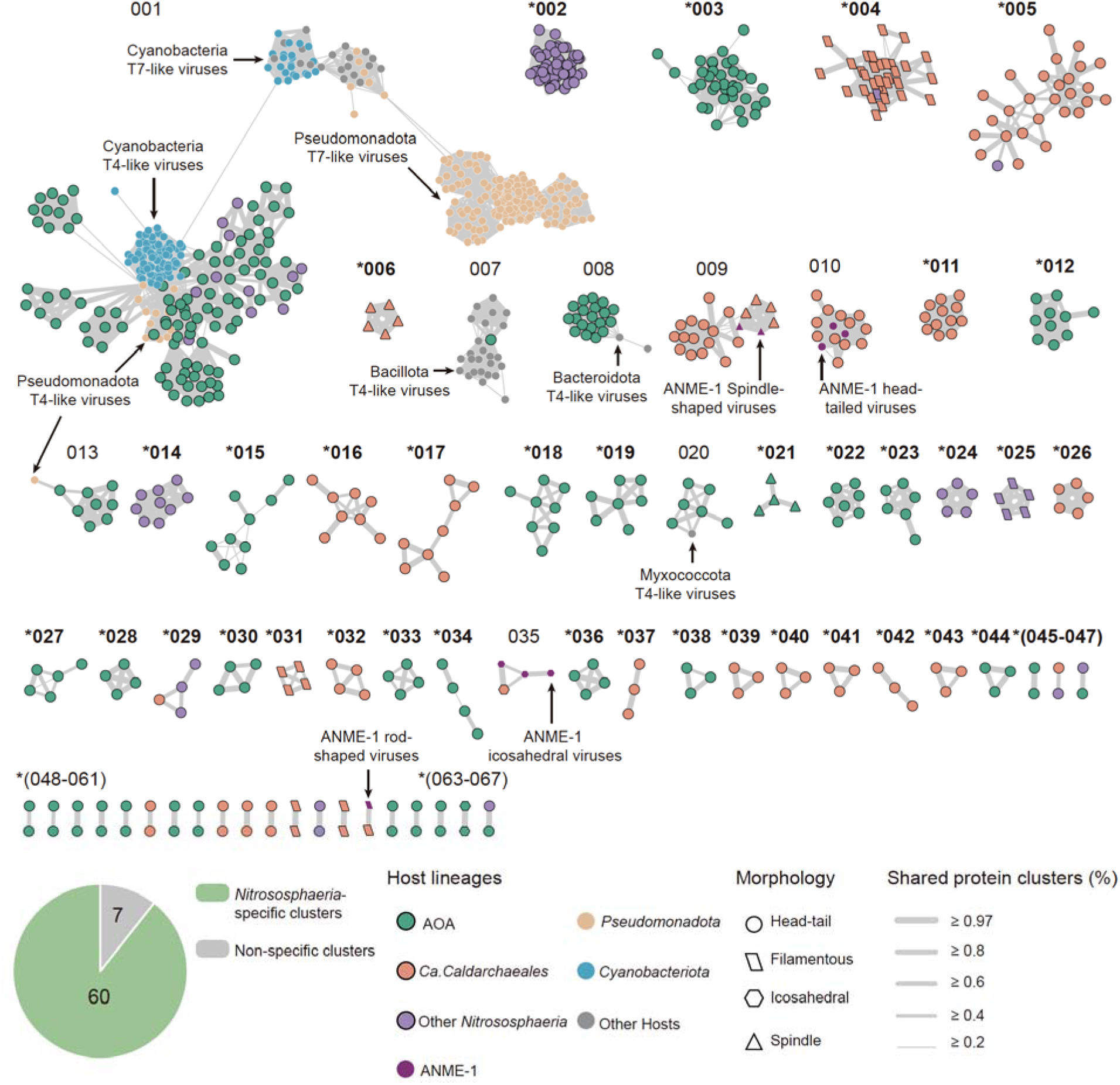
The similarity of *Nitrososphaeria*-associated viruses or mobile genetic elements (MGEs) based on their genome contents. a, Genome-content similarity network of *Nitrososphaeria*-associated viruses or MGEs within the context of the global virosphere (sequences from NCBI RefSeq v. 223). For clarity, only connected component involving *Nitrososphaeria*-associated viruses or MGEs are shown. Each node represents a viral or MGE contig and is coloured according to its host lineage. *Nitrososphaeria*-associated viruses or MGEs clustering with established viral families are annotated adjacent to their respective groups. The pie charts illustrate the number of *Nitrososphaeria*-specific viral clusters (VCs) generated from this network. The percentage of shared proteins between genomes is indicated by the thickness of the edges connecting the nodes. The shapes of the nodes denote the potential morphologies of these viruses or MGEs. Each connected component is regarded as a super viral cluster (sVC), with asterisks indicating sVCs composed exclusively of *Nitrososphaeria*-associated viruses or MGEs.

Proteome-based hierarchical clustering tree was employed to further characterize novel families based on current viral taxonomic framework ^52^. Twenty-two viral families (or family-level lineage) were found to be associated with *Nitrososphaeria*, including 14 previously undescribed ones. Except for the 22 vOTUs that were assigned to established viral families (Figs. 2, 3 and Supplementary Dataset 2), 108 vOTUs were proposed to be newly named viral families here, they include: two rod-shaped viral families (*Demeterviridae* and *Gaiaviridae*) and four head-tailed viral families (*Erebusviridae*, *Hypnosviridae*, *Morosviridae* and *Nyxviridae*), which were associated with *Ca. Caldarchaeales* and seven head-tailed family (*Apateviridae*, *Alboranviridae*, *Baleariviridae*, *Charonviridae*, *Thanatosviridae*, *Freyrviridae* and *Peleviridae*) associated with AOA (Fig. 3). In addition, nine MGE families without known capsid proteins were generated, indicating a more complex genomic diversity of the *Nitrososphaeria*-associated virome (Fig. 3 and Supplementary Dataset 2).

**Figure 3.**
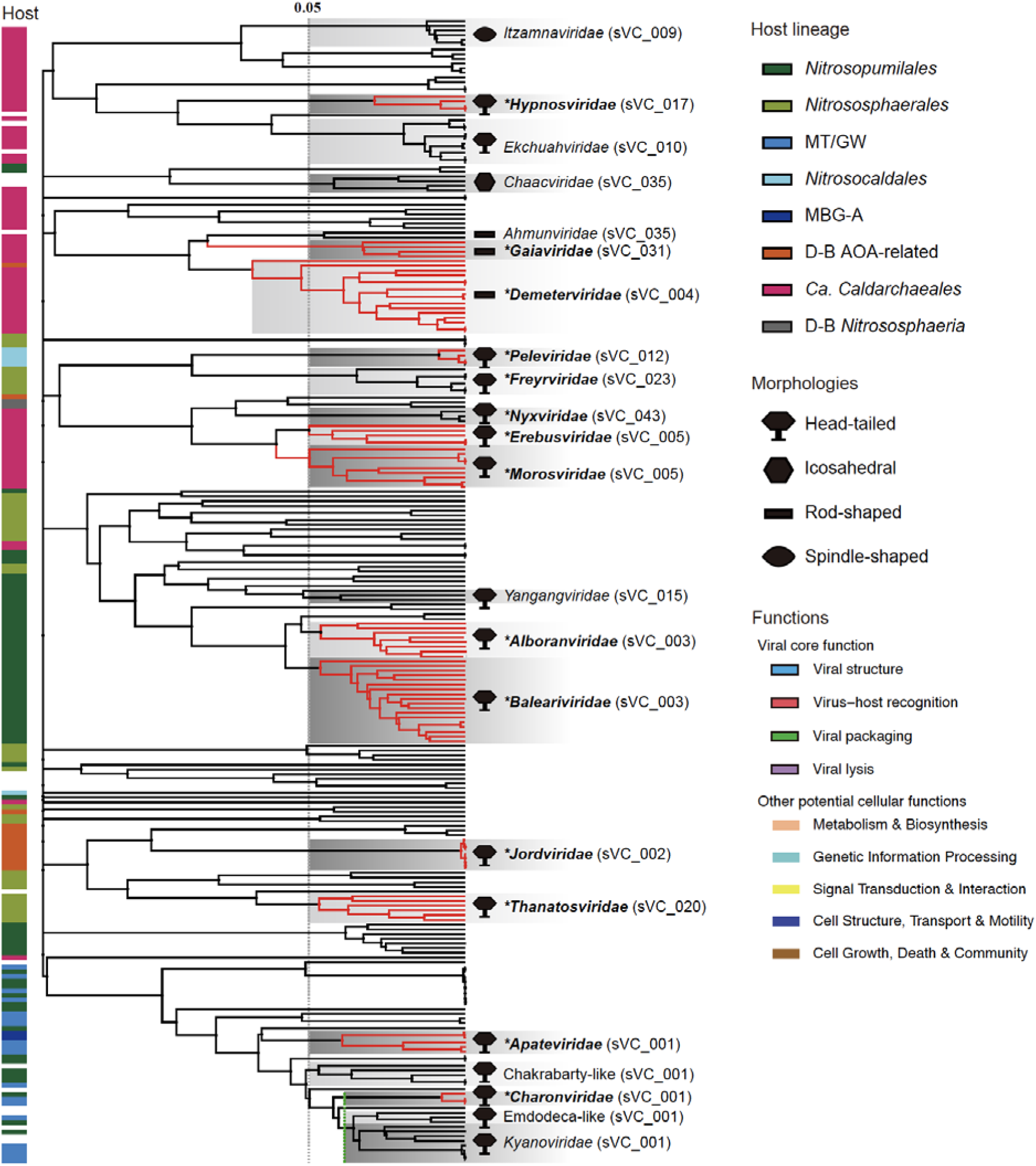
Assignments of new families within *Nitrososphaeria*-associated viruses or mobile genetic elements (MGEs). Virus proteomic hierarchy clustering tree used for the delineation of viral families. A distance threshold of 0.05 and a minimum of six shared proteins among members were used as the criteria for defining most families. *Charonviridae* were delineated by splitting from the *Kyanoviridae* clade, guided by the most recent common ancestor node. The names of the proposed families are indicated on the left side of the tree, with different virion icons representing the possible morphologies of these families. Clades corresponding to newly established viral families are highlighted on the tree with bold red branches. Tree annotation denotes host lineage and virion morphology. An asterisk is added before the names of viral families proposed in this study.

The genomes of *Nitrososphaeria*-associated viral families vary in length, and the proteins they encode exhibit diverse functional potentials (Fig. 4). *Apateviridae* contains a member with a genome length of 228 kb (vOTU_0015), which encodes 34 viral core functional genes (Supplementary Dataset 4), offering clues to the existence of previously unrecognized archaeal “jumbo” viruses ^53^. The vOTU_0015 encodes a variety of metabolic genes, notably carrying 16 Radical SAM–associated proteins, suggesting that these functions may play important roles in the survival of this virus. *Jordviridae* represent a family of temperate archaeal viruses that encode integrase and a series of Ig-like domain–containing protein, whose functions may be similar with host-recognition proteins found in other archaeal viruses ^29^. *Nyxviridae* encode TFII-like proteins (transcription factor II), a class of transcription factors typical of archaea and eukaryotes, suggesting that these viruses may hijack the archaeal transcription machinery ^28,54^. *Erebusviridae* encode MCM-like proteins (minichromosome maintenance proteins), a core DNA replication factor widely conserved in archaea and eukaryotes, suggesting that these viruses may hijack the archaeal DNA replication machinery ^55,56^. *Demeterviridae* represent a rod-shaped viral lineage encoding multiple SIRV2-like MCPs and harboring UbiG-like proteins (2-polyprenyl-3-methyl-5-hydroxy-6-methoxy-1,4-benzoquinol methyltransferase) and ArsR-like proteins, highlighting the unique quinone metabolic potential and transcriptional regulatory mechanisms of this group of archaeal rod-shaped viruses ^57^.

**Figure 4.**
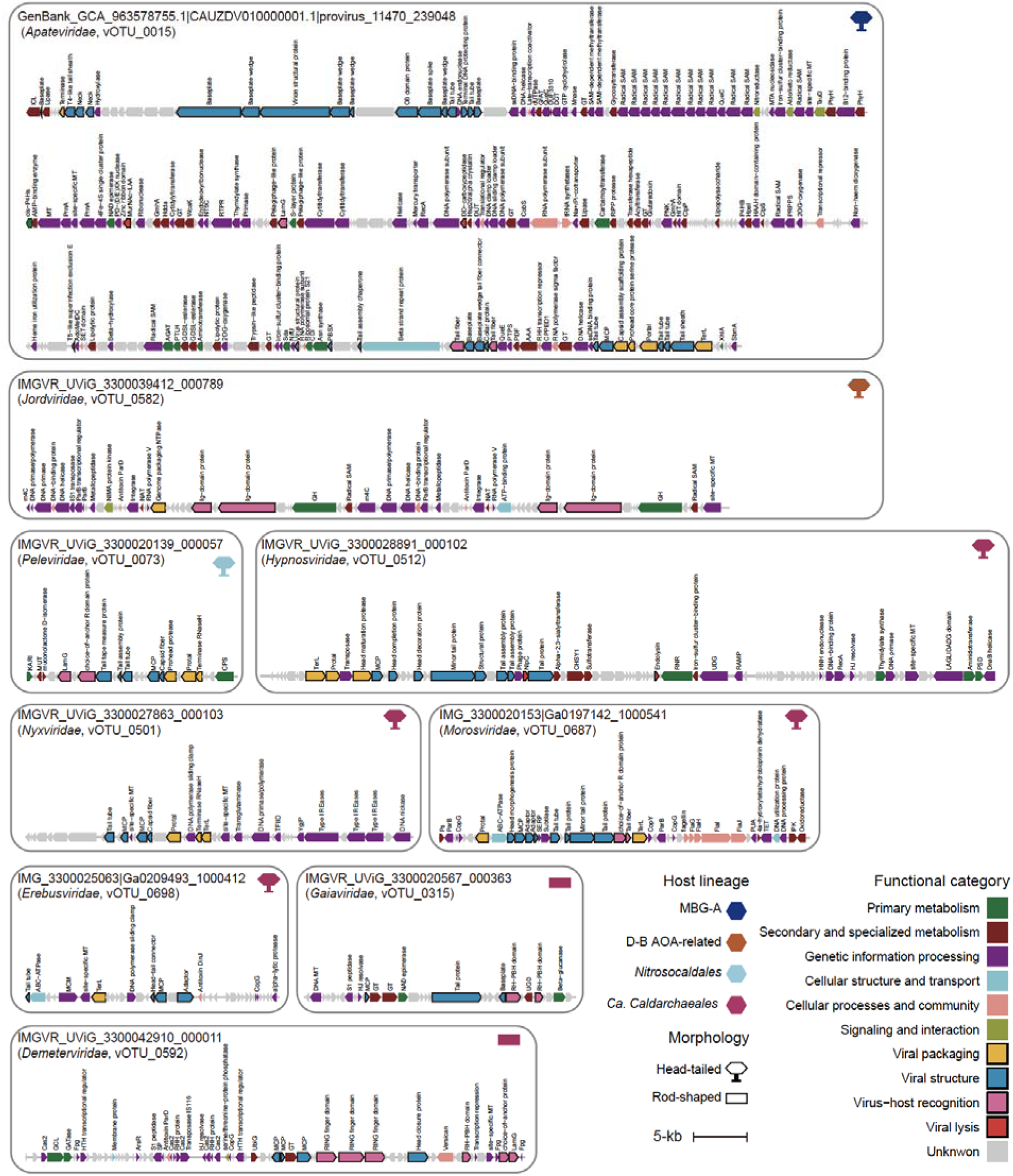
Genome organization diagrams of nine newly established viral families associated with newly expanded archaeal lineages. Arrows denote individual open reading frames (ORFs), with the colours indicating the function categories. Primary metabolism includes amino acid metabolism, carbohydrate metabolism, energy metabolism, lipid metabolism and nucleotide metabolism; secondary and specialized metabolism includes biosynthesis of other secondary metabolites, metabolism of terpenoids and polyketides, xenobiotics biodegradation and metabolism, metabolism of cofactors and vitamins and glycan biosynthesis and metabolism; genetic information processing includes replication and repair, transcription, translation, folding/sorting/degradation, information processing in viruses; cellular processes and community includes cell growth/death and cellular community. Relevant functional annotations are indicated above the corresponding ORFs. ORFs associated with viral core functions are highlighted with bold outlines. The corresponding archaeal lineages and their morphologies are indicated alongside each genome.

### The distinct evolutionary placements of *Nitrososphaeria*-associated viruses within the *Adnaviria* and *Duplodnaviria*

Bayesian (BI) and Maximum-Likelihood (ML) inference with structural modelling was used to characterize the evolution of *Nitrososphaeria*-associated viruses (Extended Data 1). Sequence-based phylogenetic inference of SIRV2-like capsomers supports a common ancestry for these rod-shaped viruses. *Sulfolobales*-associated rod-shaped viruses (*Rudiviridae*), the ANME-1–associated rod-shaped viral family (PBV300, *Ahmunviridae*), D-B-*Nitrososphaeria*-associated rod-shaped viruses (vOTU_0179), and the two *Caldarchaeales*-associated rod-shaped viral families (*Demeterviridae* and *Gaiaviridae*) each form well-supported monophyletic clades, with both posterior probabilities and maximum-likelihood bootstrap values exceeding 0.9 (90) (Fig. 5a). However, the posterior probabilities and bootstrap values supporting the nested branching relationships among these rod-shaped viral families are all below 0.7 (70), indicating that their evolutionary relationships remain unresolved. In the structural similarity network, with the exception of vOTU_0179, all SIRV2-like capsomers formed a single community (structural similarity ≥ 0.3), indicating pronounced structural similarity among them (Fig. 5b and Extended Data 2). vOTU_0179 lacks one N-terminal α-helix, whereas all other monomers possess the typical two N-terminal α-helices and three C-terminal α-helices. The monomer encoded by vOTU_0179 and one of the monomers encoded by PBV300 (GenBank accession: UYL64970.1) are highly similar in their C-terminal structures, both containing an additional eight β-sheets, suggesting that their capsids likely share a common origin. However, vOTU_0179 was not observed to encode a second SIRV2-like capsid protein as seen in PBV300, its capsomer is likely composed of *Rudiviridae*-like homodimers.

**Figure 5.**
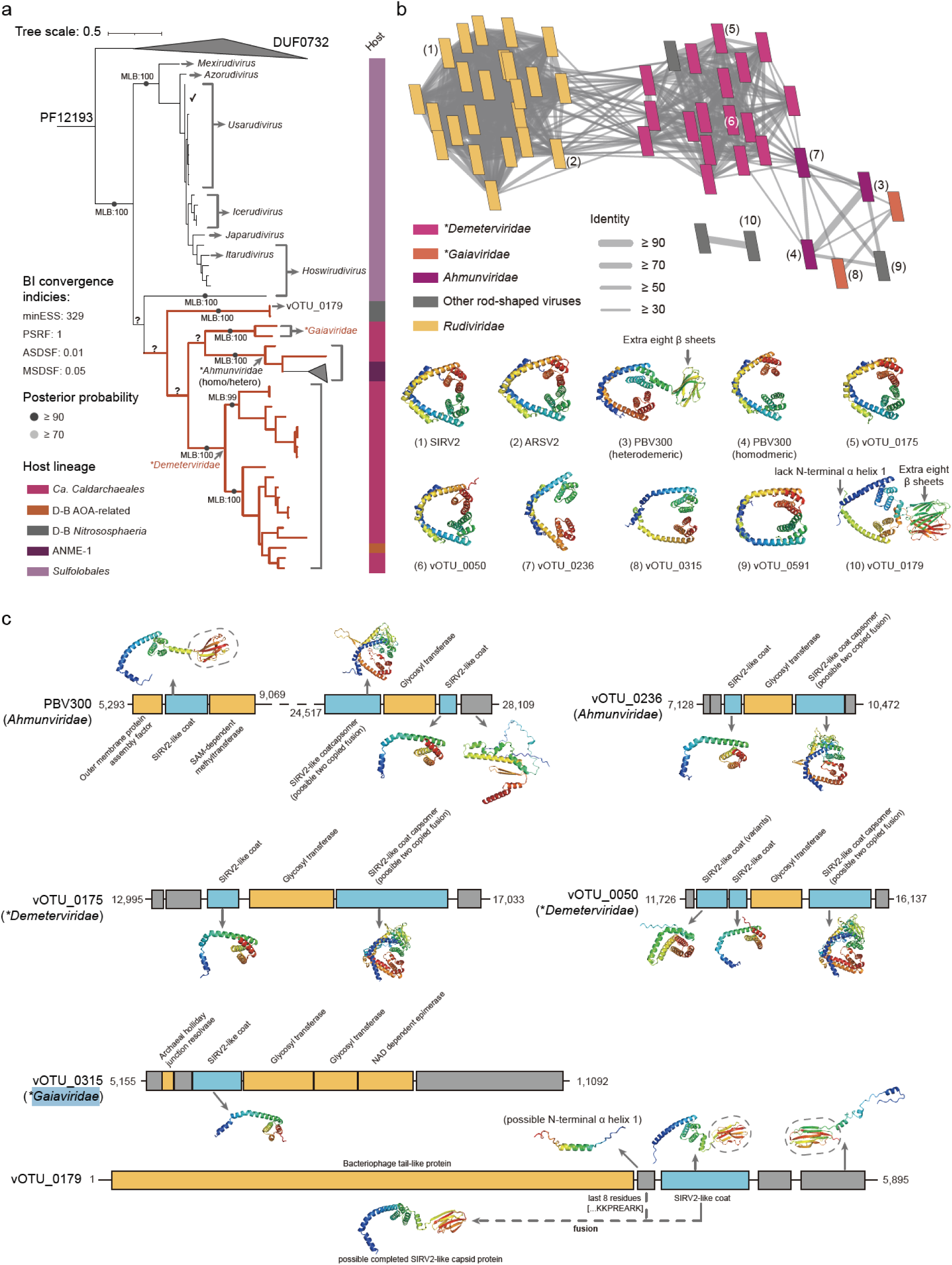
Evolution of *Nitrososphaeria*-associated rod-shaped viruses based on SIRV2-like capsomers through Bayesian inference (BI), Maximum Likelihood (ML) inference and structural modelling. a, BI tree based on the SIRV2-like capsid protein (domain accession in Pfam: PF12193). DUF732 (domain accession in Pfam: PF05305 and in VOG223: VOG14611) was used as the outgroup to root the tree. Posterior probabilities from BI are indicated on the tree branches with coloured nodes, alongside ML bootstrap values (MLB) with Blosum62+F+I+G4 as the best-fit substitutional model. The tree annotation indicates corresponding host archaeal lineages. *Nitrososphaeria*-associated rod-shaped viruses are shown as bold red branches. The names of the corresponding viral genera or families are labelled alongside the respective branches. BI convergence indices, including minimum effective sample size (minESS), potential scale reduction factor (PSRF), average standard deviation of split frequencies (ASDSF) and max standard deviation of split frequencies (MSDSF). b, Structural modelling of archaeal rod-shaped viral capsomers and their structure-based similarity network. Nodes represent viral capsomers and are color-coded according to their viral families. Edge width indicates the degree of structural similarity between capsids (only identities ≥ 0.3 are shown). Representative structures from each cluster are displayed below the networks. c, Structural modelling of proteins encoded by open reading frames (ORFs) adjacent to SIRV2-like major capsid protein (MCP) genes in *Ahmunviridae*, *Demeterviridae*, and *Gaiaviridae*. Blue boxes indicate putative structural ORFs, while orange boxes represent ORFs with other functions. Functional annotations are shown above the boxes. An asterisk is added before the names of viral families proposed in this study.

Some *Nitrososphaeria*-associated rod-shaped viruses were observed to possess unique capsid protein–encoding potential. Through structure-based functional annotation, we found that several rod-shaped viruses encode two copies of SIRV2-like capsid proteins (Fig. 5c). Similar to *Hoswirudivirus*, the two capsid protein copies are located adjacent to each other and to a glycosyltransferase gene in the genome, raising the possibility that only one of the capsid proteins has a structural function ^58^. Unexpectedly, we also identified certain rod-shaped viruses (including PBV300 and members from *Demeterviridae*) that appear to additionally encode a single gene capable of forming a complete capsomer fold (Fig. 5c and Supplementary Fig. 2), which is likewise located adjacent to a glycosyltransferase. This finding suggests the possibility that, in these viruses, a complete capsomer may be assembled from a single monomer. In addition, potential capsid protein fusion and cleavage events were observed in vOTU_0179. Although the SIRV2-like capsid protein encoded by vOTU_0179 lacks one N-terminal α-helix, fusion with the last eight residues of the adjacent gene product could generate a complete SIRV2-like capsid protein. Near the SIRV2-like capsid protein gene in vOTU_0179, a neighboring gene encodes a hypothetical protein with three N-terminal α-helices and eight C-terminal β-sheets, highly similar to the C-terminal structures of the SIRV2-like capsid proteins encoded by vOTU_0179 and PBV300. This finding suggests that SIRV2-like capsids may undergo dynamic fusion or cleavage events, exhibiting previously unrecognized functional polymorphism.

*Nitrososphaeria*-associated head-tailed viruses exhibit typical polyphyletic characteristics and generally form well-supported monophyletic clades in phylogenetic tree of major capsid protein (MCP) (Fig. 6a). The results of phylogenetic inference are consistent with the observations from the HK97-like MCP structural similarity clustering network (Fig. 6b and Extended Data 3). Eight newly identified viral families form three *Nitrososphaeria*-specific monophyletic clades, including four families associated with *Ca. Caldarchaeales* (*Morosviridae*, *Nyxviridae*, *Hypnosviridae*, and *Erebusviridae*), one family associated with *Nitrosocaldales* (*Peleviridae*), one family associated with *Nitrososphaerales* (*Freyrviridae*), two families associated with *Nitrosopumilales* (*Alboranviridae* and *Baleariviridae*), and one family associated with deep-branching AOA-related *Nitrososphaeria* (*Jordviridae*). In contrast, the remaining three AOA-related viral families (*Thanatosviridae*, *Charonviridae*, and *Apateviridae*) are phylogenetically nested with bacterial viruses, suggesting that some viruses may have undergone host switching during their evolution.

**Figure 6.**
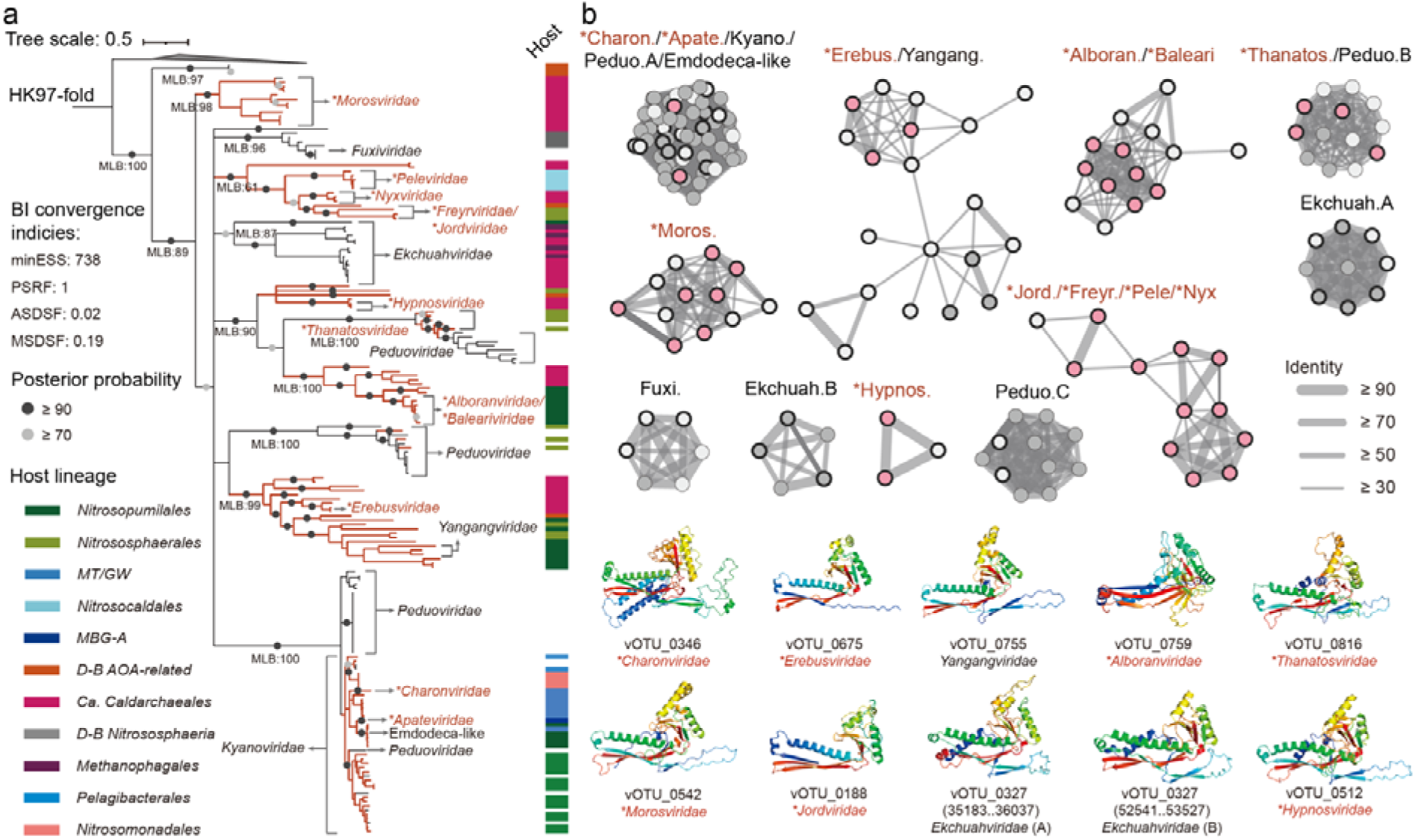
Evolution of *Nitrososphaeria*-associated rod-shaped viruses based on HK97-like major capsid proteins (MCPs) through Bayesian inference (BI), Maximum Likelihood (ML) inference and structural modelling. a, BI tree based on the HK97-like MCP sequences, rooted using archaeal dodecin (domain accession in Pfam: PF07311) as the outgroup. Because HK97-like MCPs originate from multiple distinct domains and are difficult to align accurately at the sequence level, phylogenetic inference was performed using structure-guided multiple sequence alignments (MSAs) (see Methods). Posterior probabilities from BI are indicated on the tree branches with coloured nodes, alongside ML bootstrap values (MLB) with LG+F+R5 as the best-fit substitutional model. The tree annotation indicates corresponding host archaeal lineages. *Nitrososphaeria*-associated rod-shaped viruses are shown as bold red branches. The names of the corresponding viral genera or families are labelled alongside the respective branches. BI convergence indices, including minimum effective sample size (minESS), potential scale reduction factor (PSRF), average standard deviation of split frequencies (ASDSF) and max standard deviation of split frequencies (MSDSF). b, Structural modelling of archaeal head-tailed viral HK97-like MCPs and their structure-based similarity network. Nodes represent viral HK97-like MCPs and are color-coded according to their family-level taxonomic assignments: nodes with pink indicate members of newly proposed viral families in this study, nodes with grey represent members of previously established viral families, and nodes with light grey denote members lacking family-level taxonomic assignments. Each community is labeled with the corresponding viral family name above it (with the suffix “*-viridae*” omitted). Viral families newly proposed in this study are indicated by red labels with an asterisk prefix.

### Novel auxiliary genes encoded by *Caldarchaeales*-associated rod-shaped viruses

Following the recommendations of Martin et al., potential auxiliary viral genes (AVGs) encoded by these viruses or MGEs were screened ^59^. A total of 376 auxiliary regulatory genes (AReGs), 235 auxiliary metabolic genes (AMGs) and 88 auxiliary physiological genes (APGs) were obtained (Supplementary Dataset 4). For AMGs, 2-polyprenyl-3-methyl-5-hydroxy-6-metoxy-1,4-benzoquinol methylase (UbiG) was found to be encoded by four vOTUs from the newly identified rod-shaped viral family, *Demeterviridae* (Supplementary Dataset 4). These virus-encoded UbiG-like proteins exhibit structural similarity to their cellular counterparts, all featuring the typical Rossmann-like fold (a β-sheet of seven strands surrounded by α-helices) (Fig. 7a). Molecular docking analyses showed that the virus-encoded UbiG-like proteins have affinity for 2,6-dihydroxybenzoic acid, with the benzene ring inserted into the SAM-binding pocket. Homologues of *Demeterviridae*-encoded UbiG-like protein are extremely rare in viruses, with no high-similarity homologues found in the nr database (alignment identity and coverage both ≥ 50%), highlighting its novelty. They form a distinct monophyletic clade that groups with distantly related archaeal homologues in the phylogenetic tree. This indicates their connection to archaea while retaining a clear virus-specific phylogenetic monophyly (Fig. 7a). To the best of our knowledge, CoQ is rare in archaea and there is currently no evidence that archaea and related viruses possess aromatic quinone biosynthesis pathway ^23,57^. Whether the functions of these virus-encoded UbiG-like proteins extend to the synthesis of menaquinone (MK), the major respiratory quinone in archaea ^57^, remains unknown. However, this finding provides a rare example of archaeal viruses encoding genes potentially related to quinone electron-carrier metabolism. ^60^. For AReGs, *Demeterviridae* was found to encode an ArsR-like transcriptional regulator, which was only previously reported from spindle-shaped ssDNA viruses and head-tailed viruses ^51,61^. The *Demeterviridae*-encoded ArsR-like proteins forms a well-supported, long-branched monophyletic clade with a pronounced stem branch in the phylogenetic tree and is evolutionarily distant from other bacteriophages and archaeal viruses (Fig. 7b). Regulators of archaeal filamentous or rod-shaped viruses are predominantly represented by RHH-fold proteins such as SvtR (*Rudiviridae*) and AvtR (*Lipothrixviridae*) ^62–64^. This result suggests that *Caldarchaeales*-associated rod-shaped viral genomes may employ a transcriptional regulatory mechanism similar to that of spindle-shaped viruses ^61^, providing clues to a previously unrecognized mechanism within this viral realm that awaits further validation.

**Figure 7.**
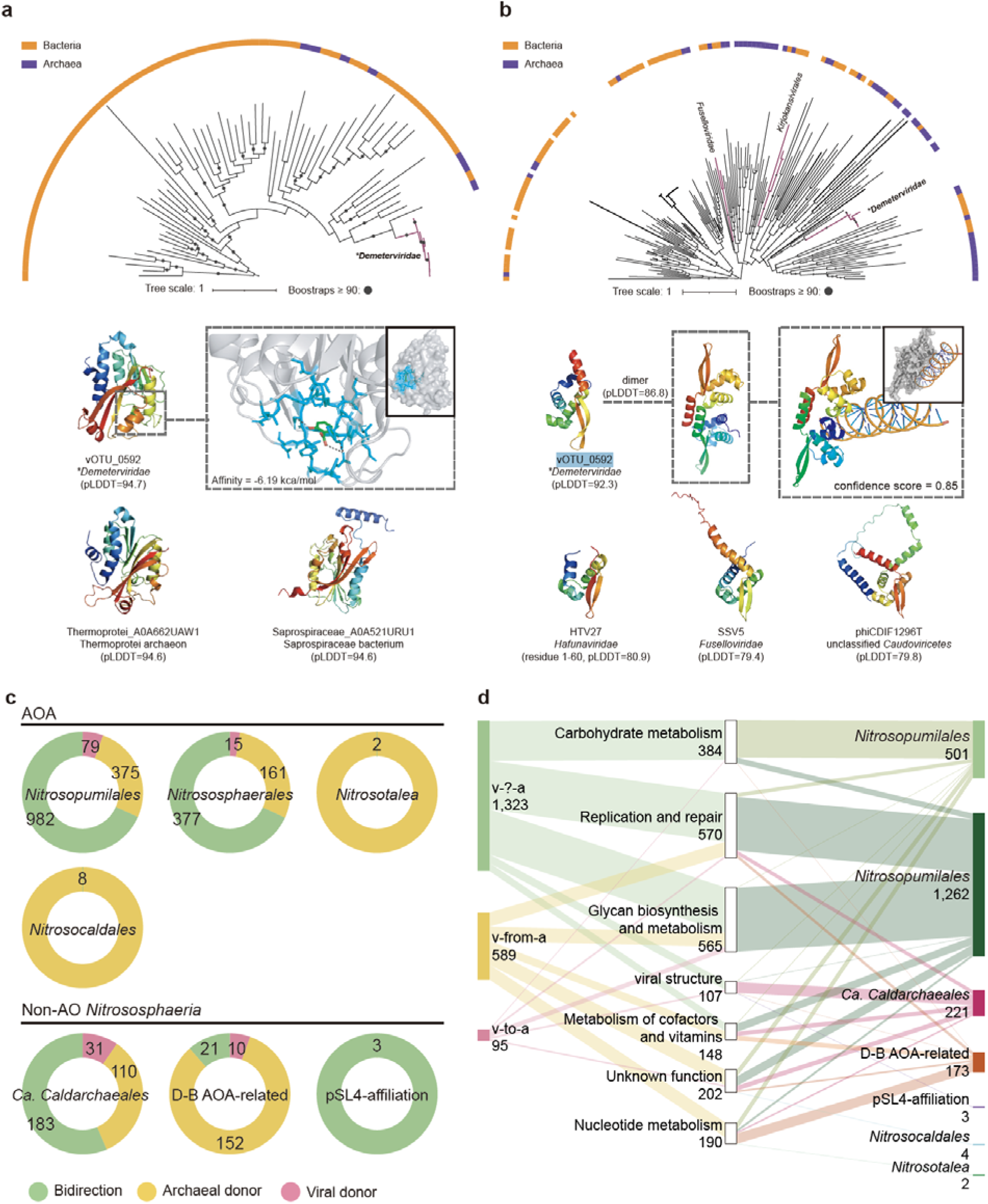
*Nitrososphaeria*-associated rod-shaped viruses encode unique auxiliary genes and exhibit a distinctive landscape of horizontal gene transfer (HGT) with their hosts. a, Maximum-likelihood (ML) phylogenetic tree of the *Demeterviridae*-encoded UbiG-like protein, with Blosum62+R10 as the best-fit model. The tree annotation includes proteins of cellular origin (from InterPro), derived from archaeal or bacterial genomes. For clarity, only clades or branches related to viruses were retained, while the others were pruned. Representative structures and their per-residue measures of local confidence (pLDDT) are displayed below the phylogenetic tree. Molecular docking simulations showed that the benzene ring of 2,6-dihydroxybenzoic acid is inserted into the Rossmann-like fold pocket of UbiG encoded by vOTU_0592. b, ML phylogenetic tree of the *Demeterviridae*-encoded ArsR-like transcriptional regulator, with LG+F+R10 as the best-fit model. Representative structures and their per-residue measures of local confidence (pLDDT) are displayed below the phylogenetic tree. Molecular docking simulations showed the binding of A-DNA to the ArsR dimer encoded by vOTU_0592. c, Proportion of HGT events with different transfer directions among various *Nitrososphaeria* lineages. Different colours of the sectors indicate the direction of HGT, and the numbers within them represent the number of observations. d, The number of HGTs with distinct directions were associated from different functional catagories and were linked with diverse *Nitrososphaeria* lineages. Bar colours represent the direction of HGT and *Nitrososphaeria* lineages, respectively. The numbers of observations were indicated beside bars.

### Viruses or MGEs serve as the primary vectors rather than donors or receptors in horizontal gene transfers

Virus–host linkages identified by spacer–protospacer matching showed lineage specificity, with no cross-lineage protospacer sharing observed (Supplementary Fig. 3a and Supplementary Dataset 5). After establishing virus-host linkages by spacer-protospacer matching, the direction of horizontal gene transfers (HGTs) mediated by viruses or MGEs were explored based on phylogenetic topologies ^65^. A total of 2,509 possible HGT events between viruses or MGEs and corresponding *Nitrososphaeria* lineages were identified (Supplementary Dataset 6 and Extended Data 4) ^65^. For AOA, 68.0% of transfers are bidirectional (n = 1,359), while 27.3% originate from archaea as donors (n = 546). Only 4.7% of transfers involve viruses or MGEs as donors (n = 94). Similarly, in *Ca. Caldarchaeales*, 56.5% of transfers are bidirectional (n = 183), 34% originate from archaea (n = 110) and 9.6% involve viruses or MGEs as donors (n = 31) (Fig. 7c). This suggests that viruses or MGEs serve as the primary vectors in these transfers, rather than donors or receptors.

Functionally, most HGTs are related to glycan biosynthesis and metabolism, replication and repair and carbohydrate metabolism, accounting for 60.5% of the total HGTs (Fig. 7d). Within glycan biosynthesis and metabolism, more than half of the HGTs (n = 283) are associated with glycosylation and are linked to *Nitrosopumilales* (Supplementary Dataset 4 and Supplementary Dataset 6). The related functional proteins include all dolichyl-phosphate-mannose–protein mannosyltransferases, glycosyl transferases (GTs), sulfatases and N-acetyl sugar amidotransferases. Since glycosylation or glycoprotein modification on the surface of archaeal viral capsids or envelopes is involved in host interactions or enhances structural stability ^66–68^, it raises the possibility that *Nitrososphaeria*-associated viruses or MGEs have incorporate host-derived functional protein modules. In addition, 87.8% of the HGTs related to viral structure were found to be associated with *Ca. Caldarchaeales*. These structural proteins include tail-related proteins and MCPs from *Erebusviridae* and *Morosviridae*, further confirming the interactions between these proposed viral families and *Ca. Caldarchaeales*.

## Discussion

In this study, we provide a deeper understanding of the diversity of *Nitrososphaeria*-associated viruses or MGEs, broadening the scope from previously reported AOA-associated viruses and identified hundreds of previously unrecognized *Caldarchaeales*-associated vOTUs for the first time. The virus–archaea associations identified in this study have not yet been validated through isolation experiments, but they provide valuable guidance for establishing future cultivation systems for *Nitrososphaeria* viruses, particularly those associated with *Ca. Caldarchaeales*. This genomic catalogue expands the known number of *Nitrososphaeria*-associated viruses or MGEs by over seven-fold ^32–36^. The mining of *Caldarchaeales*-associated viral or MGE genomes fills a critical knowledge gap of archaeal viromes. Nine previously unknown viral families were identified, including a novel rod-shaped family and an archaeal “jumbo” virus. The distinct evolutionary placement and functional characteristics of this family were elucidated through comparative genomic and phylogenetic analyses, providing new insights into the archaeal virosphere and viral evolution.

Through phylogenetic inference and structural modelling, we revealed the evolutionary relationships among archaeal rod-shaped viruses, providing new evidence for the phylogeny and taxonomy of the *Adnaviria* viral realm. In the past, only the capsomers of *Rudiviridae* were known to be composed of homodimers^64^. Here, it was found that some *Caldarchaeales*-associated rod-shaped viruses encode multiple copies of SIRV2-like MCP genes (Figs. 4 and 5c). Given the pronounced sequence homology between the MCPs encoded by *Demeterviridae* and those of *Rudiviridae*, and the fact that no *Rudiviridae* with homodimeric capsomers have yet been identified ^30,64,69^, it is possible that in *Demeterviridae* only one MCP copy is functional, while the other may have been inactivated through natural selection ^70–72^. *Rudiviridae*-like viruses with a single functional MCP subunit likely represent a more ancestral state, whereas viruses encoding two related MCPs evolved through duplication and divergence ^73^. The observation of a fusion gene encoding two SIRV2-like MCPs further advances this hypothesis: the ancestral capsomer of rod-shaped viruses may have been composed entirely of a single protein, which subsequently gave rise to homodimers and, later, heterodimers during evolution. Through structural modelling of the MCP encoded by vOTU_0179 and its adjacent genes, we found that this MCP appears to be undergoing ongoing cleavage, potentially generating additional protein products (Fig. 5c). Because its structure is incomplete, this MCP may no longer be functional. The MCP encoded by vOTU_0179 is structurally highly similar to one of the MCPs encoded by *Ahmunviridae* PBV300 (GenBank accession: UYL64970.1). PBV300 may retain functionality in only a single MCP (GenBank accession: UYL64989.1), thereby forming a homodimeric capsomer. However, because PBV300 also encodes a fusion gene containing two MCP subunits (GenBank accession: UYL64987.1), the bona fide capsid architecture of these rod-shaped viruses remains unknown. Taken together, our evidence indicates that *Nitrososphaeria*-associated rod-shaped viruses, *Rudiviridae*, and *Ahmunviridae* are highly similar in the evolutionary and structural characteristics of their capsid proteins. In addition, comparative genomics analyses revealed pronounced genomic synteny among these rod-shaped viruses (Supplementary Fig. 4). This raises the need to reconsider whether the placement of *Ahmunviridae* within *Maximonvirales* as a lineage parallel to *Ligamenvirales* is justified, or whether *Ahmunviridae* should instead be incorporated into *Ligamenvirales*.

Viruses have been found to encode diverse AVGs that manipulate host metabolism, thereby creating favourable conditions for completing their life cycles and indirectly contributing to global biogeochemical cycles ^74–77^. In archaea-associated viruses, such as those infecting methanogenic archaea, Asgard superphylum, halophilic archaea, virus-encoded metabolic genes have been identified that potentially modulate archaeal methane-oxidization, sulphur metabolism, nucleotide and amino acid metabolisms ^27,78,79^. Regarding AOA-associated viruses or MGEs, previous studies have discovered that viruses may manipulate host ammonia oxidation metabolism by carrying the *amoC* gene and participate in host vitamin synthesis ^34,35^. Through the exploration of functional repertoires for *Nitrososphaeria*-associated viruses, we provide new insights into the functional mechanisms of archaeal viruses. An important example is provided by viral ArsR-like and UbiG-like protein, which represent a rare AReG and AMG, respectively, encoded by a previously unrecognized rod-shaped viral family. If the viral ArsR-like protein is functional in the archaeal virocell, it might be able to hijack the arsenic response of *Ca. Caldarchaeales*, thereby facilitating viral replication within the host cell. Considering that the habitat of *Ca. Caldarchaeales* is rich in metals and metalloids (including arsenic) and exhibits highly active redox dynamics^24,80,81^, this process may be essential for the completion of the *Demeterviridae* life cycle. Phylogenetically, *Demeterviridae*-encoded ArsR falls on a long-branch, virus-specific clade supports their viral origin, rather than recent horizontal gene transferring from archaea.

The more unusual finding is the viral carriage of UbiG-like protein, an enzyme implicated in CoQ biosynthesis. Quinones (e.g. ubiquinone, menaquinone) are key electron carriers in membrane redox chains, yet they are considered rare in archaea and corresponding viruses ^23,57,82^. The detection of *Demeterviridae*-encoded UbiG-like protein suggests a potential viral contribution to the redox balance of archaeal membranes, such as altering the local pool of quinone electron carriers ^57,83^. They might influence electron flux, respiration efficiency, reactive oxygen species (ROS) levels, or even redox signaling ^83–85^. In hyperthermal habitats of *Ca. Caldarchaeales*, control of archaeal quinone pools may confer a fitness advantage to related viruses by maintaining a more favorable intracellular environment for replication. On the other hand, the *Demeterviridae*-encoded UbiG-like protein, together with other methyltransferases encoded by archaeal viruses, may perform similar functions in counteracting host defense mechanisms, representing an exaptation of archaeal gene functions by viruses ^86^. Since *Ca. Caldarchaeales* has not been observed to encode UbiG-like proteins, the origin of these genes in *Demeterviridae* remains unknown. Overall, the discovery of *Demeterviridae* not only expands the boundaries of the archaeal virosphere and offers new insights into viral evolution, but also provides direction in how viruses might adapt to extreme environments in terms of transcriptional regulation and host metabolic reprogramming.

Our results reinforce the view that viruses or MGEs act predominantly as gene vectors. This supports the “virus world” model, considering viruses or other selfish genetic elements as intermediaries in gene exchange rather than as main sources of new genes ^87^. The prevalence of bidirectional HGTs highlights reciprocal coevolution between viruses or MGEs and archaeal hosts, evoking the arms-race dynamics observed in archaeal virus–host systems ^88,89^. The functional enrichment of glycan biosynthesis and glycosylation among transferred genes implies that viruses may co-opt host surface modification machinery to refine capsid stability or host recognition—a phenomenon similarly observed in eukaryotic viruses adapting envelope glycosylation ^90,91^. It is hypothesized that the frequent HGT events between archaea and viruses may have driven the diversification of archaeal envelope biochemistry and the emergence of niche specialization in ammonia oxidizing lineages.

## Methods

### Assembly of *Nitrososphaeria*-associated CRISPR database

The *Nitrososphaeria* phylum comprises several major clades: AOA, other AOA-related lineages and *Ca. Caldarchaeales*. To produce a comprehensive genomic dataset, reference genomes were retrieved from GenBank and IMG/M ^41,42^, encompassing a total of 2,137 genomic assemblages. Specifically, 1,830 assemblages (comprising 322,784 genomic contigs) were obtained from GenBank; 307 assemblages (containing 23,395 genomic contigs) were sourced from IMG/M.

To identify CRISPR spacers within these genomes, similar to previous studies ^27,28^, CRASS v.0.3.8 and MinCED v.0.4.2 (CRISPR recognition tool, CRT) were employed using the following parameters to detect spacers and discard those with low-complexity and short length ^92,93^: word size = 7, spacer length = 26 and a minimum of three directed repeats (DRs). This step resulted in the identification of 37,485 *Nitrososphaeria*-associated spacers, which were subsequently used for the detection of viruses or MGEs and downstream analyses.

The metagenome-derived spacers that are associated with *Nitrososphaeria* were detected to further expand the diversity of these CRISPR spacers. First, a total of 170 metagenome-derived assemblages (including 58 from the Malaspina expedition and 22 from our previous studies) from diverse ecosystems were retrieved and downloaded from IMG/M and ENA (European Nucleotide Archive) (Supplementary Dataset 1)^42,45^. These metagenomes were from 23 different studies, focusing on extreme natural habitats and including 92,037,046 assembled contigs. These include four studies from deep-sea, three two from sea ice, four studies from a hypersaline environment and one study from an acidic environment. Second, the CRISPR spacers found in these contigs were identified using the same approach described above, producing 1,828,641 spacers and 1,918,441 DRs. Third, all DRs were aligned to *Nitrososphaeria* genomes using MMseqs2 with parameters ^94^: word size = 7, pairwise alignment identity = 1, percentage coverage of DRs = 1, mismatches = 1. This step produced 196,551 spacers derived from 170 metagenomes. Finally, this process generated a *Nitrososphaeria*-associated spacer database containing 234,036 spacer sequences (Supplementary Fig. 1 and Extended Data 5).

As in previous studies ^28^, CRISPR-Cas cassette prediction and subtyping for archaeal contigs and genomes were conducted using CRISPRcasIdentifier v1.0.0 ^95^, employing a combination of Classification and Regression Trees (CART), Support Vector Machine (SVM) and Extremely Randomized Trees (ERT) algorithms for classification and regression. The analysis was performed with the following parameters: –mode ERT SVM CART, -hmm HMM1 HMM2 HMM3 HMM4 HMM5. Subtype assignment was determined through a consensus-based approach, where only subtype classifications consistently predicted by all three models (CART, SVM and ERT) were accepted. The final subtype was selected based on the highest mean probability and the lowest standard deviation among the predictions (Supplementary Fig. 3b and Supplementary Dataset 7).

### Mining and filtering *Nitrososphaeria*-associated viruses or MGEs from IMG/VR (v.4)

The IMG/VR (v.4) database, contigs from 170 metagenomes and archaeal genomes served as the primary resources for mining *Nitrososphaeria*-associated viral or MGE genomes. At first, the archaeal viral or MGE mining pipeline was validated using CRISPR spacers derived from archaeal genomes and host predictions from the IMG/VR (v.4) database and other studies to minimize false positives ^33–36,38,96,97^. The 37,485 CRISPR spacers from *Nitrososphaeria* genomes were aligned to IMG/VR (v.4) database using MMseqs2 with the parameters: word size = 7, pairwise alignment identity = 0.9, percentage coverage of spacers = 0.9, mismatches = 1, resulting in 368 targeted. Among these results, the hosts of 46 viruses or MGEs were also predicted to be *Nitrososphaeria* in IMG/VR (v.4). Due to some viral contigs being assigned to non-*Nitrososphaeria* hosts in IMG/VR (v.4) metadata, the genome-content similarity between viruses or MGEs and hosts was applied to reduce the false positives: i) the open reading frames (ORFs) of these viral contigs were predicted by Prodigal v.2.6.3 and aligned to the NR database (2024.05.20) with 1e-5 as the E-value cut-off ^98^; ii) the *Nitrososphaeria*-related hits were counted and normalized based on viral contig lengths; iii) for each genome, it was observed that at least 0.2 ORFs per 1-kb genome could be matched to *Nitrososphaeria* (Supplementary Dataset 2); iv) further validation was conducted to assess the applicability of this screening criterion to previously reported *Nitrososphaeria*-associated viruses or MGEs:

1. the 87 identified *Nitrososphaeria*-associated viral or MGE genomes were downloaded from GenBank or provided by data owner;
2. the 46 representative genomes were generated from 48 *Nitrososphaeria*-associated viral genomes through MMseqs2 with parameters ^94^: pairwise alignment identity = 0.95, pairwise alignment coverage for shorter sequences = 0.9;
3. the 83 out of 85 viral genomes derived from GenBank meet this criterion. Two isolated spindle-shaped viruses (NSV4 and NSV5) failed to pass the test ^37^. This indicates that the approach is also applicable to most previously reported *Nitrososphaeria*-associated viral or MGE genomes.

After validating the filtering approach, it was used as a cut-off for viral-host genome-content similarity to further filter the results obtained from spacer-protospacer matching. By applying this approach, 171 *Nitrososphaeria*-associated viruses or MGEs were filtered from 368 spacer-targeted viral or MGE candidates. Similar to previous studies ^29,99^, all relevant archaeal CRISPR-Cas systems were evaluated using CRISPRCasFinder with default parameters ^100^. Among the identified CRISPR arrays, 91.2% (n = 301) exhibited an evidence level of 4, characterized by a repeat conservation index exceeding 70% and spacer identity below 8% (Supplementary Dataset 2). The remaining 29 viruses or MGEs, corresponding to evidence levels below 4 for CRISPR arrays, were retained for two reasons: 1) they passed genome-content similarity filtering; 2) in genome-wide analyses, they clustered with other viruses or MGEs possessing CRISPR arrays with an evidence level of 4.

In addition, IMG/VR (v.4) contains 1,457 viruses or MGEs that were predicted to be associated with *Nitrososphaeria*, although these contigs were not retrieved through spacer-protospacer matching. The 1,457 viruses or MGEs predicted to be associated with *Nitrososphaeria* in IMG/VR (v.4) were tested. First, 699 representative contigs were clustered using MMSeq2 with parameters: pairwise alignment identity = 0.9, pairwise alignment coverage for shorter sequences = 0.9. Second, these contigs were tested by genome-content similarity calculation, resulting in 441 *Nitrososphaeria*-associated viruses or MGEs. Finally, 612 *Nitrososphaeria*-associated viruses or MGEs were identified and filtered from IMG/VR (v.4).The completeness of these viral or MGEs were evaluate using CheckV ^101^.

### Expanded mining of *Nitrososphaeria*-associated viruses or MGEs from 170 metagenomes

The *Nitrososphaeria*-associated CRISPR spacer database was aligned to from 170 metagenomes using MMseqs2 with the same approach and parameters ^94^. Contigs containing DRs and those shorter than 2-kb were removed before performing alignment. This resulted in 1,951 contigs matched by *Nitrososphaeria*-associated CRISPR spacers. To remove duplications, 1,951 contigs were aligned to previously identified *Nitrosospaherota*-associated viruses or MGEs set using MMseqs2 with same parameters same parameters ^94^. This step generated 1,895 contigs, including 1,396 representative contigs clustered using the same approach described before. To recall as many as possible viruses or MGEs, these contigs were used to filter *Nitrososphaeria*-associated viruses or MGEs using three multiple viral identification pipelines, including geNomad, VirSorter2 and DeepVirFinder^102–104^. Multiple pipelines were employed for virus identification based on the following considerations: 1) geNomad demonstrates superior overall classification performance compared to other pipelines ^103^; 2) VirSorter2 (VS2) is recommended when integrating multiple pipelines ^105^; 3) DeepVirFinder (DVF), utilizing an alignment-free approach, is well-suited for identifying unique viral sequences ^102^; 4) the identification results from each pipeline complement one another, enhancing recall while mitigating cellular contamination in the dataset^106–108^. For each pipeline, precision was prioritized over sensitivity; therefore, stringent virus detection strategies were adopted to minimize potential false positives: 1) geNomad was used to detect viral or MGE contigs using conserved mode (“--conservative”), resulting in a more strict set of viruses with high confidence; 2) VS2 was used to detect viral contigs under high-confidence mode (’--high-confidence-only’), retaining results with classification score ≥ 0.9 and classification score ≤ 0.7 with viral hallmark ≥ 1; 3) The DVF used to detect viral contig with parameters: score ≥ 0.9 and p-value ≤ 0.01. To validate this hybrid viral detection pipeline, 20% of sequences were randomly selected from previously identified *Nitrososphaeria*-associated viruses or MGE and were used as a test set to evaluate the efficiency of the approach. Based on the test, this viral or MGE identification approach achieved a 74% recruitment rate, which was considered acceptable for further viral or MGE identification (Supplementary Dataset 8).

Using the validated hybrid viral identification approach, 419 viral or MGE contigs were identified from 1,396 metagenome-derived contigs. These contigs had already been targeted by *Nitrososphaeria*-associated CRISPR spacers in previous steps. The viral-host genome-content similarity was used to filter these contigs as described before, resulting 108 *Nitrososphaeria*-associated viral or MGEs. Relevant archaeal CRISPR-Cas systems of these contigs were evaluated using CRISPRCasFinder with default parameters ^100^. The completeness of these viral or MGEs was evaluated using CheckV ^101^.

### Detection of viruses or MGEs integrated in *Nitrososphaeria* genomes

Proviruses integrated into *Nitrososphaeria* genomes were identified using the “find-proviruses” program in geNomad ^103^. Integrases were first detected with the MMseqs2 plugin in geNomad ^94^, applying an E-value cut-off of 1e-5. Additional parameters were set as follows: a minimum gene-level score of 0.4 for flagging proviral genes using the conditional random field model; a minimum total virus marker score of 12 for proviruses lacking integrases or not located at scaffold edges; a minimum total virus marker score of 8 for proviruses encoding integrases or located at scaffold edges; a maximum allowed distance of 10-kb between provirus boundaries and integrases for boundary extension; and a maximum allowed distance of 5-kb between provirus boundaries and tRNAs for boundary extension. This process resulted in the identification of 30 *Nitrososphaeria*-associated proviruses.

To remove redundancy, the 30 proviral or MGE genomes performed self-alignment and were compared against previously identified *Nitrososphaeria*-associated viral or MGE genomes using the same approach and parameters described above. This filtering step yielded 24 non-redundant *Nitrososphaeria*-associated proviruses (Supplementary Dataset 2).

### Host assignments of *Nitrososphaeria*-associated viruses or MGEs

The host assignments of *Nitrososphaeria*-associated viruses or MGEs were primarily based on: 1) CRISPR-spacers-associated host genomes; 2) information provided by previous studies; 3) genome-content similarity between viruses or MGEs and hosts. For *Nitrososphaeria*-associated viruses or MGEs that match CRISPR spacers from multiple distinct hosts, similar with previous study ^29^, the host of a virus or MGE targeted by CRISPR spacers was assigned based on the taxonomy of the genome from which the spacer with the highest identity originated. For *Nitrososphaeria*-associated viruses or MGEs that were identified based on oligonucleotide pattern similarity, the most possible host genomes associated with these viruses or MGEs were determined through alignment against *Nitrososphaeria* genomes using MMseqs2 with the parameters: 1e-5 as the E-value, iteratively searched two times ^94^. A host genome was linked to a viral or MGE genome if they have the most shared ORFs. This approach was also applied in IMG/VR (v.4) ^96,109^. For *Nitrososphaeria*-associated viruses or MGEs reported in previous studies, some host information was retrieved from related literature. For those viruses or MGEs with uncertain host information ^35^, the related viral ORFs were subject to MMseqs2 against all *Nitrososphaeria* genomes with the same approach. The 35 Marthaviruses derived from López-Pérez et al. (PRJNA484324) have been authorized for use and are included in this study ^34^. The archaeal host phylogenetic tree was calculated based on 46 concatenated ribosomal proteins by IQ-Tree2 (Supplementary Note 2)^110^. The archaeal lineages were determined based on their phylogenetic placements and previous studies^2,8,21,22,24,111,112^.

### Comparative genomics and family-/species-level demarcation of *Nitrososphaeria*-associated viruses or MGEs

For species-level unit assignments, viral or MGE species (vOTUs or MGE OTUs) were defined using 95% nucleotide identity threshold over 90% of the full length using MMseqs2. For family-level unit assignments, similar to previous studies ^27,29,31^, new families were assigned based on the genome-content similarity network and the genome-wide proteomic tree. Using DNA virus sequences longer than 2-kb from NCBI RefSeq v.223 as reference genomes, vConTACT2 was applied (with default parameters) to compute the genome-content-based similarity network of those *Nitrososphaeria*-associated viruses or MGEs. *Nitrososphaeria*-associated viral or MGE genome pairs that shared at least 20% protein clusters and encoded at least six genes were filtered from genome-content-based similarity network, generating a subset of network, which was used to guide taxonomic assignments. In the network, viruses not related to *Nitrososphaeria*-associated viruses or MGEs were discarded to achieve a clear layout. Genome-wide proteomic tree of those genomes included in the subset of network were calculated by ViPTree v.4. The viruses or MGEs were assigned to the same family if they were represented by a cohesive group in the genome-content-based similarity network and a monophyletic group in genome-wide proteomic tree (with threshold 0.05) ^29^. The network and tree were visualized by Cytoscape v.3.10.2 and iToL v.6, respectively ^113,114^. The nomenclature of the proposed viral family was referred to Greek or Norse mythology (Supplementary Note 3).

### Annotation of viral or MGEs genomes

Viral or MGE species (vOTUs or MGE OTUs) were defined using 95% nucleotide identity threshold over 90% of the full length using MMseqs2. The family-level taxonomic assignments of *Nitrososphaeria*-associated viruses or MGEs based on reference taxonomic database were assigned using 1) VITAP with custom database: this database includes all viral reference genomes collected in ICTV VMR-MSL40, high-quality vOTUs with genus-level taxonomy collected in IMG/VR (v.4) and all viral genomes collected in NCBI RefSeq v.223 ^96,115,116^; 2) CAT with NCBI RefSeq v.223 database ^116,117^; 3) geNomad with default parameters ^103^. The CAT only assigned class/family level units for 10 contigs; the geNomad and VITAP assigned class/family level units for 683 and 676 contigs, respectively. Hence, the taxonomic assignments were mainly based on geNomad and VITAP. If geNomad or VITAP showed consistency at higher taxonomic levels (Phylum or Class), but one of the pipelines produced annotations at lower taxonomic levels, then the results with the lower taxonomic level annotations were used. The results with discrepancies between VITAP and geNomad were compared against the NCBI RefSeq v.223 using MMseqs2 with default parameters and subsequently subjected to manual verification. The consensus VCs generated from vConTACT2 were used to expanded and furtherly curated the viral taxonomic assignment ^118^. The members of each proposed family were furtherly expanded based on consensus vOTU.

ORFs encoded in *Nitrososphaeria*-associated viruses or MGEs were predicted by Prodigal v.2.6.3 with metagenome mode and annotated using the KEGG Ortholog database (January, 2025) ^98^, Pfam-A v.35, COG2020, Swiss-Prot database (January, 2025), VOG223 database and NR database (August, 2025), and HHpred ^119–122^. Potential AVGs encoded by these viruses or MGEs with length over 10-kb were screened based on proposal from Martin et al.^59^. The functional screening criteria excluded all genes associated with viral core functions (replication, recombination, repair, transcription, translation, packaging and lysis), nucleotide metabolism and one-carbon pool processes, nucleic acid modification and immune evasion, structural and host recognition functions (including all CAZymes and polysaccharide synthesis related functions), dNTP supply, as well as various proteases or multifunctional enzymes. In addition, candidate genes were filtered by examining whether viral genes were present in their flanking regions. Considering the incompleteness of viral contigs, once an AVG was confirmed, its homologues were also considered as AVGs, even if no viral genes were present in the flanking regions. The maximum-likelihood phylogenetic trees of ArsR and UbiG were constructed. For ArsR, protein sequences from archaea, bacteria, bacteriophages and archaeal viruses were retrieved from InterPro (PF01022). After merging these with the ArsR protein sequences encoded by rod-shaped viruses, CD-HIT v. 4.8 was used to cluster the sequences from different sources (at 0.5 pairwise identity with global alignment). Seqkit v.2.5 was then applied to randomly sample 100 sequences from each category ^123^, or to include all sequences if fewer than 100 were available. Multiple sequence alignment (MSA) was performed with MUSCLE5 ^124^ and trimAl v.1.4 was used to remove sites with more than 30% gaps ^125^. Maximum-likelihood phylogenetic inference of the MSAs was carried out with IQ-TREE2 ^110^, with 1,000 bootstraps and LG+F+R10 as the best-fit substitution model (Extended Data 8). For UbiG, protein sequences from archaea, bacteria and eukaryotes were retrieved from InterPro (IPR004951). These were merged with rod-shaped virus-encoded UbiG-like sequences and maximum-likelihood phylogenetic inference was performed using a similar procedure, with Blosum62+R10 as the best-fit substitution model (Extended Data 6). The maximum-likelihood phylogenetic trees were visualized with iTOL v.6 ^114^.

### Determining of *Nitrososphaeria*-associated viral or MGE horizontal gene transfer events

HGT events between *Nitrososphaeria*-associated viruses (vOTUs) and their archaeal hosts were identified following an established pipeline ^65^. After removal of proviral sequences, viral and archaeal sequences were aligned and shared protein families (PFs) were identified. Potential contamination was rigorously screened based on lineage, GC content, scaffold length and genomic context, retaining only robust alignments (Supplementary Note 4 and Supplementary Dataset 9).

Filtered protein families underwent phylogenetic analysis using MUSCLE5 alignment, trimAl v.1.4 trimming and IQ-TREE2 tree construction with the LG+F+R5 model. Trees were rooted via minimal ancestral deviation and potential HGT events were classified based on the phylogenetic relationships between viral and archaeal sequences (Extended Data 4). Transfer direction labels (“v-from-a”, “v-to-a”, “v-?-a” and “non-HGT”) were assigned following predefined criteria considering sister and cousin clade compositions. Iterative screening was performed with temporary reassignment of labels to ensure accurate HGT detection across multiple rounds of analysis (Supplementary Note 4).

### Phylogenetic inference of *Nitrososphaeria*-associated viruses or MGEs

To investigate the evolutionary relationships among rod-shaped and head-tailed viruses associated with *Nitrososphaeria* within the virosphere context, phylogenetic analyses were performed based on key viral proteins, including the SIRV2-like MCPs and HK97-like MCPs. For rod-shaped viruses, SIRV2-like MCP sequences were aligned with viral genomes of *Adnaviria* realm from GenBank using MMSeqs2 (E-value ≤ 1e-5). Some viruses within *Rudiviridae* (*Hoswirudivirus*) encoded adjacent duplicate MCP subunits, proteins adjacent to these SIRV2-like MCPs were annotated using HHpred to determine whether additional distantly related SIRV2-like MCPs are present. SIRV2-like MCPs directly adjacent to glycosyl transferases were used to generate homodimer sequences, aligned using MUSCLE5 (ensemble mode) to generate MSAs. Sites with more than 30% gaps were removed using trimAl v.1.4 ^125^. The trimmed MSAs were subsequently used for BI (300 million generations) and ML (3000 bootstraps) phylogenetic inference based on MrBayes and IQ-TREE2, respectively ^110,126^. The DUF732 domain-containing protein (PF05305) with three α helices (similar with C-terminal α helices of SIRV2-like MCP), identified via FoldSeek structural alignment as the closest homolog ^127^, served as an outgroup for rooting the rod-shaped viral MCP phylogeny.

HK97-like MCP sequences exhibited significant diversity, encompassing multiple families from Pfam-A v.35, CDD, and VOG223 ^120,128,129^. Thus, a structure-guided approach was employed to perform phylogenetic inference of HK97-like MCPs. First, putative HK97-like MCP sequences were collected through domain annotation. Second, structural modelling of these sequences was performed using AlphaFold2 implemented via ColabFold. Third, all MCPs were aligned under the guidance of structural information using Reseek and MUSCLE5 to generate a structure-based multiple sequence alignment (MSA)^130^. Fourth, the resulting MSA was used to build a hidden HMM with HMMER3 ^131^, which was then applied to search for homologs among all *Nitrososphaeria*-associated viral proteins (E-value ≤ 1e-5). Fifth, based on all identified *Nitrososphaeria*-associated viral HK97-like MCPs, BLASTp searches were conducted against RefSeq v.223 to retrieve viral homologs using thresholds of 0.5 for both pairwise coverage and sequence identity. Sixth, all newly identified HK97-like MCP sequences, together with archaeal Dodecin sequences as an outgroup ^86^, were subjected to a second round of structural modelling using AlphaFold2, and a second structure-based MSA was constructed following the same procedure. Finally, this MSA was used for phylogenetic inference using the same methods.

### Structure modelling and molecular docking for viral proteins

All structural modelling was performed using AlphaFold2 implemented via ColabFold with default parameters. The models with the highest pLDDT scores were selected as the best-predicted structures. Foldseek was used to perform all-vs-all structural alignments for SIRV2-like MCPs and HK97-like MCPs separately (E-value ≤ 1e-5; identity ≤ 0.3) ^127^. The resulting alignments were visualized as networks in Cytoscape ^113^, using bitscore as the edge weight. For molecular docking of UbiG, the following steps were applied: 1) structural data of a potential substrate containing an aromatic ring and phenolic hydroxyl groups (2,6-dihydroxybenzoic acid) were retrieved from PubChem ^132^; 2) docking simulations were performed using HDOCK ^133^, with the search space covering the entire β-sheet core region; and 3) docking results were visualized using PyMOL ^134^. For molecular docking of ArsR, the following steps were conducted: 1) ArsR monomers were assembled into the biologically active dimeric form; 2) A-DNA was generated from a random genomic fragment of *Demeterviridae* (vOTU_0592) using Web 3DNA 2.0; 3) docking simulations were performed using HDOCK, with the search space covering the entire cluster of C-terminal α-helices; and 4) docking results were visualized using PyMOL.

## Data availability

All nucleotide and protein sequences of viral or mobile genetic elements, spacers from the CRISPR database, phylogenetic trees, structural prediction files are available in FigShare (https://figshare.com/s/a2e94d686028b674a905). The related metagenomes used in this study can be accessed via the URL provided in Supplementary Dataset 1 through IMG/M. Related genomes from IMG/VR database is available at https://img.jgi.doe.gov/cgi-bin/vr/main.cgi. Other information and metadata are provided as Supplementary Datasets.

## Code availability

No custom code was generated and used.

## Acknowledgements

We thank Jilu Han for his help with the installation and debugging of the software used in this study. We thank the support of the high-performance servers of the Center for High Performance Computing and System Simulation, Laoshan Laboratory (Qingdao), the High-Performance Biological Supercomputing Center at the Ocean University of China, the Marine Big Data Center of the Institute for Advanced Ocean Study of the Ocean University of China and the IEMB-1, a high-performance computing cluster operated by the Institute of Evolution and Marine Biodiversity. This study was supported by the National Natural Science Foundation of China (No. 42120104006, 42176111 and 42306111), the Ocean Negative Carbon Emissions (ONCE), 2024 Graduate Self-directed Research Project (2024ZZKY, 202461036) and the Fundamental Research Funds for the Central Universities (202172002, 201812002, 202461036 andrew McMinn).

## Author contributions

Conceptualization: K.Z. and Y.L.; Analysis: K.Z. and H.Y.; Database: D.P.-E.; Writing manuscript: K.Z.; Review and editing: Y.L., A.M., F.W. and M.W.; Computational resource: D.P.-E., M.L-P., S.L., G.W.N., C.H. and M.W.; Funding acquisition and supervision: Y.L., A.M. and M.W.

