## Supplementary Note for "A global viromic survey reveals unprecedented viral diversity within ammonium-oxidizing archaea and related lineages"

### Supplementary Notes

#### Supplementary Note 1 Detailed pipeline for mining *Nitrososphaeria*-associated viruses or MGEs

The construction of the *Nitrososphaeria*-associated viruses or MGEs catalogue involved detailed mining and filtering steps. Initially, 170 metagenomes from diverse extreme habitats were screened for viral sequences using a hybrid viral identification approach (Supplementary Dataset 1) ^1-16^. A CRISPR spacer database containing 234,036 spacers (66,890 non-redundant spacers) derived from *Nitrososphaeria* genomes and environmental metagenomes was established (Extended Data 6). Viral sequences were identified through alignment of spacers against viral genomic datasets (including IMG/VR v.4 and metagenome-derived viruses) ^17^. Subsequently, candidate sequences underwent stringent filtering using protein-sharing analyses to reduce false positives.

#### Supplementary Note 2 *Nitrososphaeria* phylogenetic inference based on ribosomal proteins

Gene prediction for 346,179 genome contigs derived from 2,137 *Nitrososphaeria* assemblages was performed using Prodigal v.2.6.3. Ribosomal protein-associated sequences were identified and extracted based on the COG2020 database, encompassing a total of 87 ribosomal protein clusters (ribosomal COGs). These ribosomal COGs were aligned individually using MUSCLE5 with the '-super5' mode ^18^, and subsequently, hidden Markov models (HMMs) were constructed using HMMER3 ^19^. The predicted open reading frames (ORFs) from the *Nitrososphaeria* genomes were aligned against the ribosomal COG HMMs through 'hmmsearch', employing an E-value threshold of 1e-10 for both sequence and domain alignments. For host phylogenetic inference, related *Nitrososphaeria* genomes and *Thermoprotei* reference genomes were used to calculate maximum likelihood tree, with a subset of 46 concatenated ribosomal proteins. Specifically, aligned regions corresponding to these ribosomal proteins were extracted from the ORFs of each assemblage based on alignment information obtained via the '-A' flag in 'hmmsearch'. The extracted ribosomal COG regions were re-aligned using MUSCLE5 with default parameters. Multiple sequence alignments (MSAs) were refined using trimAl v.1.4 to remove fast-evolving positions, specifically columns containing over 10% gaps. A set of 60 *Thermoprotei* assemblages obtained from RefSeq v. 223 were included as an outgroup in the phylogenetic analyses. The final concatenated alignment, comprising 5,963 amino acid positions from the 46 ribosomal proteins, was used to infer a maximum-likelihood phylogeny employing IQ-Tree2 under the LG+C60+F model, with 1,000 bootstrap replicates, following methodological recommendations from Spang et al. Taxonomic assignments of the *Nitrososphaeria* groups were performed based on previous consensus phylogenetic placements of archaeal lineages from previous studies ^20-25^.

#### Supplementary Note 3 Detailed description of nomenclature for newly proposed *Nitrososphaeria*-associated viral families

The nomenclature of the newly proposed *Nitrososphaeria*-associated viral families follows a unified thematic framework, in which the family suffix “-viridae” is combined with prefixes derived from Greek mythology, Norse mythology, or the names of regions within the Mediterranean Sea. For viral families associated with *Nitrosopumilales*, several names were derived from either Greek mythology or Mediterranean geography. *Charonviridae* is named after Charon, the ferryman of Greek mythology who transports souls across the rivers separating the living world from the underworld. Three viral families belonging to the “Marthavirus” group were named after distinct regions of the Mediterranean Sea. These include *Alboranviridae*, derived from the Alboran Sea located between the Iberian Peninsula and North Africa, and *Baleariviridae*, named after the Balearic Sea in the western Mediterranean. For viral families associated with *Nitrososphaerales*, names were selected primarily from Norse and Greek mythology. *Freyrviridae* is named after Freyr, a Norse god associated with fertility, prosperity, and favorable environmental conditions. *Thanatosviridae* derives its name from Thanatos, the Greek god of death. For viruses associated with *Nitrosocaldales*, the family *Peleviridae* was named after Pele, the Polynesian goddess of fire and volcanoes. Several viral families infect hosts from mixed or uncultivated *Nitrososphaeria*-related groups. *Apateviridae* is named after Apate, the Greek goddess of deception. For viral families associated with deep-branched AOA-related lineages (D-B AOA-related), two names rooted in mythology were adopted. *Jordviridae* is derived from Jörð, the Norse personification of the Earth. *Demeterviridae* is named after Demeter, the Greek goddess of agriculture, and was chosen in consideration of its sister-group relationship with *Ahmunviridae*, whose name originates from a Mayan god of agriculture. This parallel nomenclature reflects their close phylogenetic relationship. Another rod-shaped viral family associated with *Ca. Caldarchaeales*, *Gaiaviridae*, deriving its name from Gaia, the primordial Greek deity of the Earth.

#### Supplementary Note 4 Detailed description of viral or MGE horizontal gene transfer events filtering

The *Nitrososphaeria*-associated vOTUs and corresponding host's genomic contigs (with length over 250-kb) were used to determine the horizontal gene transfer events using similar approach with previous study^26^. First, the proviral regions were removed from host’s genomic contigs. The vOTUs were aligned to corresponding archaeal contigs using MMseqs2 with parameters: pairwise alignment identity = 0.95, percentage coverage of viral/MGE contigs = 0.5. Second, the pairwise alignment scores (bitscores) were used to cluster protein families (PFs) using Markov Cluster Algorithm (MCL) with inflation 2 ^27^. PFs containing both viral or MGE and archaeal representatives were retained. Only pairwise alignments from the same PFs were retained. Third, all related proteins were assessed for their potential contamination using four scores: 1) lineages of related archaeal genomes per viruses or MGEs gene (+1 for inter lineages; +3 for intra lineages); 2) GC content within one standard deviation of the median genome (+1 otherwise 0); 3) archaeal genomic contigs was greater than half of the scaffold N50 of related assemblages; 4) scores_t-based taxonomy of related genomic context (upstream and downstream 2.5 kbp) of per related ORF (+1 for archaea; 0 for viruses/MGEs; -1 for bacteria and eukaryotes). Only alignments with summed scores of over 2 were retained as non-contaminated pairs (Supplementary Dataset 10). Fourth, all filtered viral-host PFs were aligned using MUSCLE5 using default parameters ^18^. The fast-evolving sites (possessing over 20% gaps) were removed from alignment columns by trimAl v.1.4 ^28^. Alignments with at least 50 amino acid positions were retained. Fifth, all multiple sequence alignments (MSAs) of PFs were performed phylogenetic inference by IQ-TREE2 with LG+F+R5 substitution model and SH-aLRT statistical support calculation. All trees were rooted using a minimal ancestral deviation approach. In final, the HGT candidates were screened out based on following criteria as described in previous study ^26^:

1. Starting from the first viral/MGE leaf node, identify its sister clade (i.e., the most closely related phylogenetic clade) and cousin clade (i.e., the second most closely related phylogenetic clade), and prune them into a subtree (without altering the original topology). All clades refer not only to those containing more than two clusters but also to branches consisting of a single leaf.

2. For the pruned subtree, if the cousin clade of the viral or MGE leaf is composed entirely of archaea, label the viral leaf as "v-from-a (recipient)" and record the labels of all archaeal leaves with the shortest branch length (if multiple leaves share the shortest branch length, include all) within the cousin clade as "archaeal donor member."

3. If the sister clade of the viral or MGE leaf contains archaeal leaves, but the cousin clade consists entirely of viruses/MGEs, label the viral/MGE leaf as "v-to-a (donor)" and record the labels of the archaeal leaves with the shortest branch length (if multiple leaves share the shortest branch length, include all) within the sister clade as "archaeal recipient member."

4. If the cousin clade of the viral leaf contains both viral/MGE and archaea, label the viral/MGE leaf as "v-?-a" and record the current viral/MGE leaf's label, as well as the labels of the viral and archaeal leaves in its cousin and sister clades as "Unknown transfer direction."

5. If the sister clade of the viral or MGE leaf is entirely viral, but the cousin clade is entirely archaeal, label all related viral leaves as "non-HGT."

6. During the traversal process, when a viral or MGE leaf's sister clade and cousin clade are pruned to form a subtree, if this subtree contains other viral or MGE and archaeal leaves that have already been defined, assign new attributes to these leaves before proceeding with steps 1-5: viral leaves defined as "v-from-a" are re-labelled as archaeal, viral/MGE leaves defined as "v-to-a" retain their viral or MGE label, archaeal leaves defined as "archaeal donor member" retain their archaeal label, and archaeal leaves defined as "archaeal recipient member" are re-labelled as viral.

7. Assign a temporary label of "viral/MGE " to all members of the archaeal recipients and a temporary label of "archaeal" to all members of v-from-a. Then, re-ran the HGT detection to examine whether members that were defined as non-HGT during the first round of HGT screening can be assigned HGT-related attributes (such as v-to-a, v-from-a, v-?-a, archaeal recipient, or archaeal donor). This process was iterated three times.

### Supplementary Figure Legends

Figure 1. The workflow for mining, filtering, and identifying viruses or MGEs primarily included the prediction of archaeal CRISPR systems, viruses or MGEs mining, genome filtering, and the definition of viral or MGE operational taxonomic units (vOTUs or MGE OTUs). Only genomes encoding detectable capsid or related proteins were considered viruses; others were classified as MGEs.

Figure 2. Sequence and structural modeling of a fused open reading frame encoding SIRV2-like coat capsomers (vOTU_0175). The two SIRV2-like capsid domains within the sequence are highlighted in red and yellow, respectively, and displayed in the 3D structure.

Figure 3. Overview of *Nitrososphaeria* CRSPR-Cas systems and their linkages with viruses or mobile genetic elements (MGE). a, Spacer-sharing network among *Nitrososphaeria* genomes. Nodes represent archaeal genomes and are colored by distinct archaeal lineages. An edge is generated between two genomes if they share spacer sequences, suggesting that their CRISPR systems may target the same viral or MGE species. Bold edges indicate that the shared spacer sequences have corresponding protospacers present in the *Nitrososphaeria*-associated viruses or MGEs catalogue generated in this study. b, Examples of CRISPR-Cas systems harboured in *Nitrososphaeria* genomes/contigs targeting viruses/mobile genetic elements (MGEs) from diverse taxonomic groups. Each CRISPR-Cas cassette is annotated with the number of direct repeats (R) and spacers (S), genomic/contig coordinate, and the type of CRISPR-Cas system. The IDs of the corresponding archaeal genomes/contigs are shown below each cell schematic. The morphologies of the targeted viruses/MGEs are illustrated using distinct virion schematics, with the associated sequence IDs.

Figure 4. Comparative genomics–based gene synteny diagram of rod-shaped viruses possibly belonging to *Ligamenvirales*, including *Rudiviridae*, *Ahmunviridae*, *Gaiaviridae*, and *Demeterviridae*. The colors of the linkages between open reading frames indicate pairwise sequence identity based on PSI-BLASTp.

### Supplementary Dataset

Dataset 1. The metagenomic assembled contigs used in this study were derived from public databases, encompassing metagenomes originating from four types of extreme environments: acidic, deep sea, hydrothermal, and sea ice habitats. The associated metadata includes study names, IMG accession numbers, accessible URLs, and corresponding publications.

Dataset 2. The metadata of *Nitrososphaeria*-associated virus and mobile genetic element (MGE) contigs. This table encompasses contig IDs, associated metagenome/genome identifiers, geographic information, related research project names, taxonomic information, viral classifier confidence scores, genomic features, data source, environmental origin details, host information, and the proportion of shared genes between each virus (or MGE) and archaeal lineages.

Dataset 3. Genome-content similarity between *Nitrososphaeria*-associated virus and mobile genetic element (MGE) contigs and other reference viral genomes.

Dataset 4. Functional annotation of proteins encoded by *Nitrososphaeria*-associated viruses and mobile genetic elements (MGEs). This table includes the IDs of *Nitrososphaeria*-associated viruses and MGEs, family-level taxonomic information, viral cluster assignments generated by vConTACT2, contig lengths, putative morphotype information, archaeal lineage information, environmental context, protein IDs, functional domain accession numbers, functional domain annotations, viral auxiliary gene labels and corresponding database names.

Dataset 5. Sharing of CRISPR spacer sequences among archaeal genomes. This table includes genome IDs of the paired archaeal genomes, their taxonomic information, and the viral or mobile genetic element (MGE) contigs targeted by the corresponding spacers.

Dataset 6. Horizontal gene transfer (HGT) events between viruses or mobile genetic elements (MGEs) and their associated archaeal lineages. This table includes open reading frame (ORF) pairs derived from viruses or MGEs and archaea, the inferred direction of transfer, taxonomic information of both archaeal and associated viral lineages, functional annotations and accession numbers, enzymatic reaction equations, and metabolic functional classifications across three distinct aspects.

Dataset 7. CRISPR cassettes identified in the archaeal genomes involved in this study were typed using CRISPR-Cas-Identifier. This table summarizes the results generated by CRISPR-Cas-Identifier, including archaeal genome IDs, cassette types, cassette coordinates, and the associated Cas-related proteins.

Dataset 8. Performance of viral recall by three different viral identification pipelines based on a test set of viruses or mobile genetic elements (MGEs). VS2 and DVF denote VirSorter2 and DeepVirFinder, respectively.

Dataset 9. Scoring matrix for identifying horizontal gene transfer (HGT) events based on the approach proposed by Irwin et al. Specifically, potential HGTs are quantitatively evaluated across four distinct criteria: transfer conditions (cross-lineage or intra-lineage), G+C content, scaffold N50, and the taxonomic information of the open reading frames (ORFs) flanking the candidate HGT region.

### Extended Data

Data 1. The Bayesian and Maximum-likelihood tree of major capsid proteins encoded by *Nitrososphaeria*-associated viruses.

Data 2. The structural predication of SIRV2-like capsid proteins from AlphaFold2.

Data 3. The structural predication of HK97-like capsid proteins from AlphaFold2.

Data 4. The *Nitrososphaeria*-associated CRISPR spacers sequences, which were predicted from MinCED and CRASS.

Data 5. The Maximum-likelihood tree of proteins that were potentially involved in horizontal gene transfers (HGTs). These trees were used to determine HGT events based on their topologies.

Data 6. The Maximum-likelihood tree of UbiG and ArsR proteins encoded by *Nitrososphaeria*-associated viruses and cellular organisms.

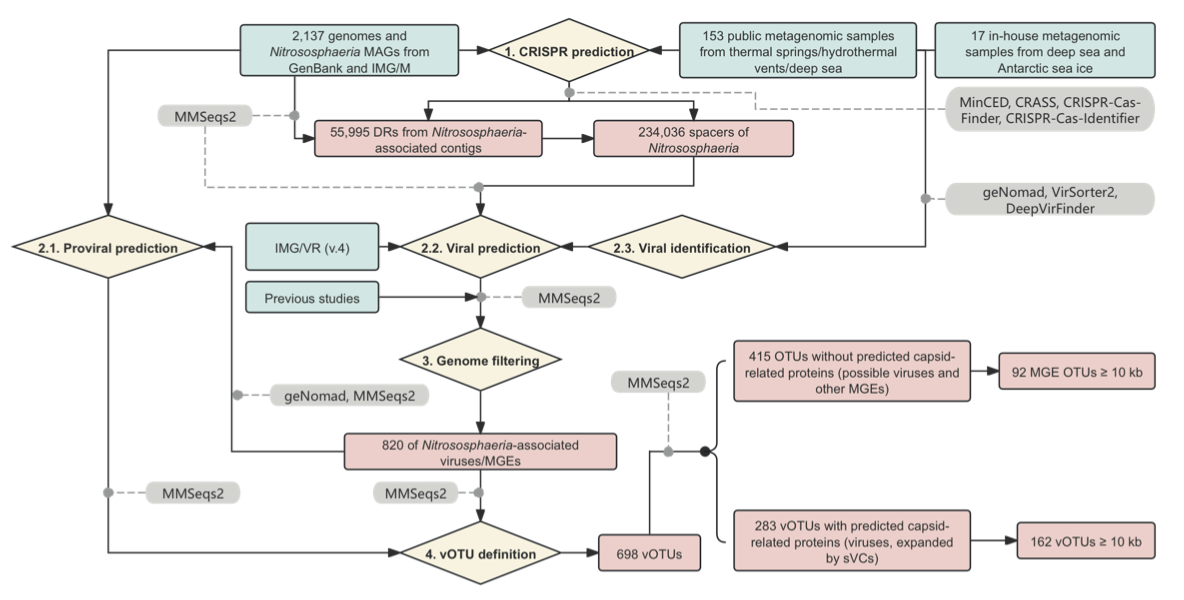

Supplementary Figure 1

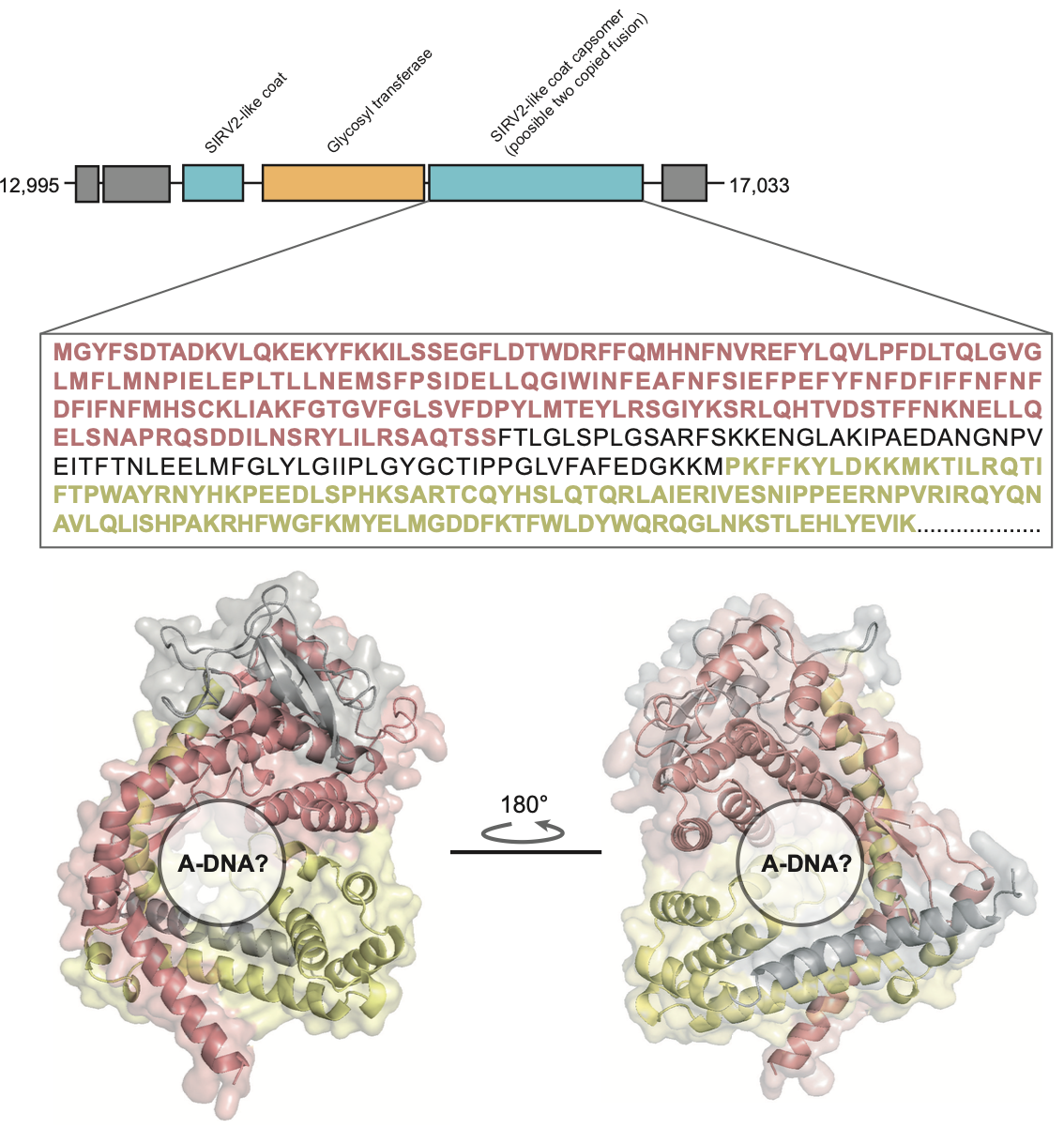

Supplementary Figure 2

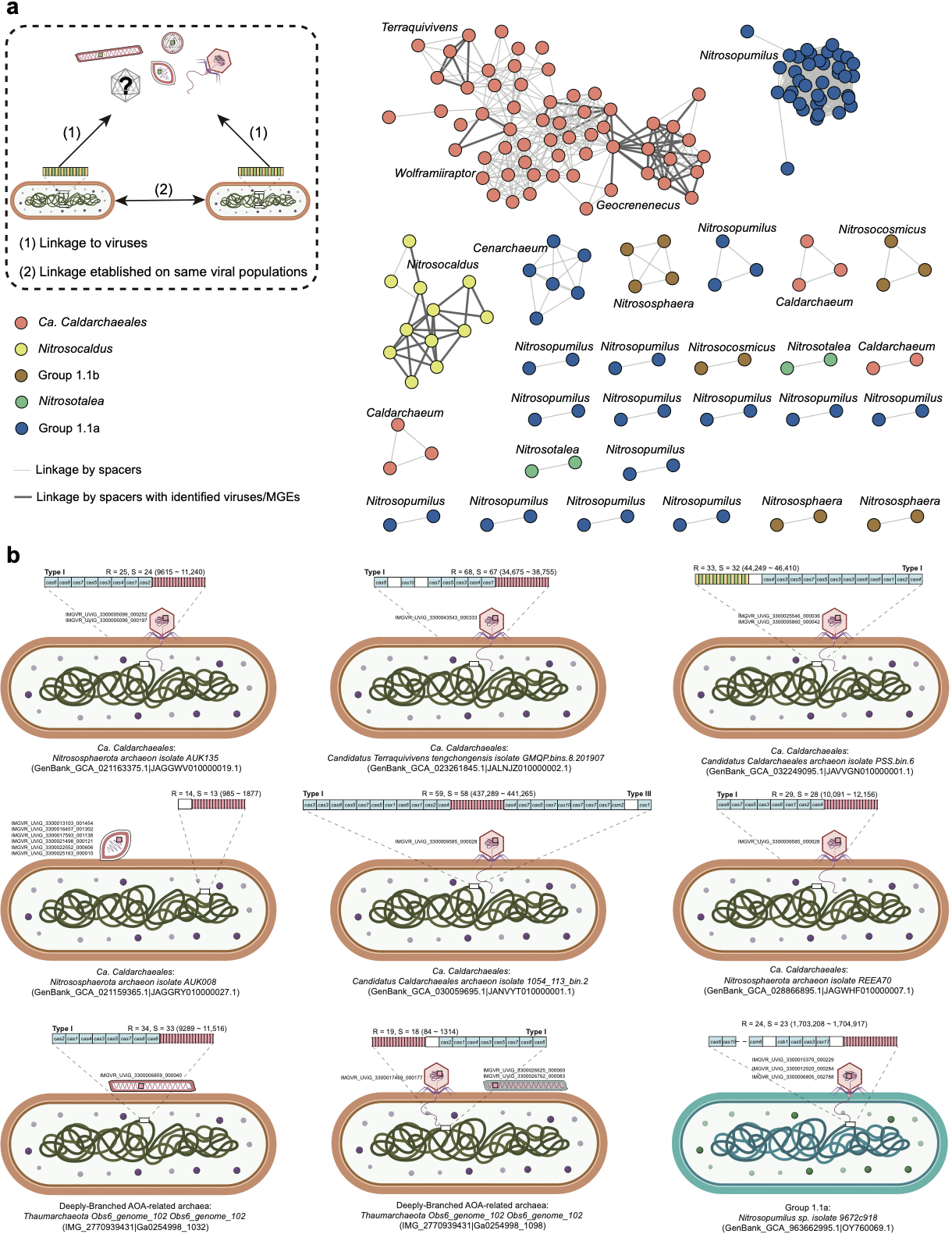

Supplementary Figure 3

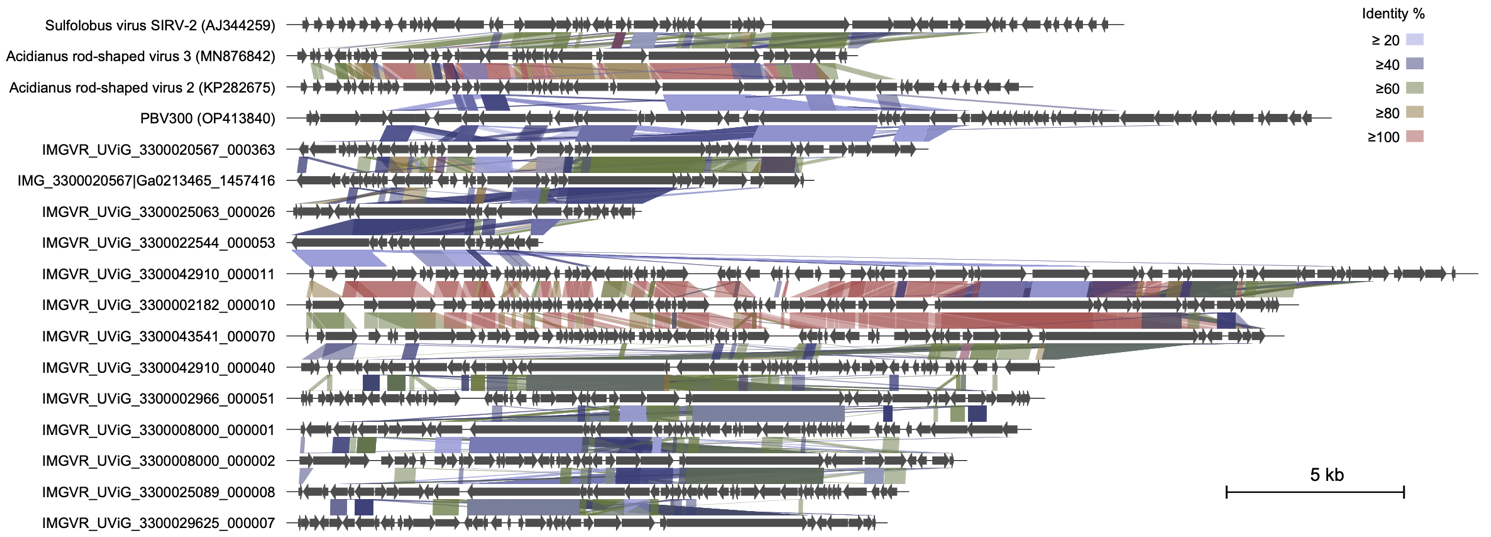

Supplementary Figure 4
